# Evolutionarily guided transcription factor design programs novel T cell states

**DOI:** 10.1101/2024.11.06.622344

**Authors:** Oliver Takacsi-Nagy, Austin Hartman, Andy Y. Chen, Yajie Yin, Gabriella C. Reeder, Courtney Kernick, Johnathan Lu, Alison K. McClellan, Colin J. Raposo, Nicole E. Theberath, Patrick K. Yan, Justin Eyquem, Theodore L. Roth, Ansuman T. Satpathy

## Abstract

Protein-coding genes in the human genome evolved via modular rearrangement of domains from ancestral genes^1^. Here, we develop a scalable, evolutionarily guided method to assemble novel protein-coding genes from constituent domains within a protein family, termed DESynR (Domain Engineered via Synthesis and Recombination) genes. Using primary human chimeric antigen receptor T cells as a model system, we find that the expression of DESynR Activator Protein-1 (AP-1) transcription factors (TFs) significantly outperforms the overexpression of natural AP-1 TFs in multiple functional assays *in vitro* and *in vivo*. Top DESynR AP-1 TFs exhibit non-intuitive architectures of constituent domains, including from TFs that are not canonically expressed in T cells. DESynR AP-1 TFs induce broad transcriptional and epigenetic reprogramming of T cells and, in some cases, lead to the development of non-natural T cell states, engaging gene expression modules from disparate human cell types. Taken together, we demonstrate that novel configurations of existing protein domains may uncover non-evolved genes that program cell states with therapeutically relevant functions.

## Main

The sequencing of the human genome revealed that the evolution of new human protein-coding genes relies primarily on repurposing protein domains from ancestral genes, rather than *de novo* sequence creation^1^. This is borne out in the expansion of protein families, where new genes are derived from novel linear rearrangements of domains via addition, deletion, or shuffling^1–7^. These observations demonstrate that evolving new functions does not necessarily require new sequences; rather, new functions can emerge from existing sequences placed in new genetic contexts. While methods for directed evolution via mutagenesis or CRISPR-editing enable screening of base pair-level sequence evolution for improved protein or cellular functions, similar technologies for domain-level evolution are comparatively underdeveloped^8,9^. Specifically, domain shuffling methods have largely focused on optimizing or innovating the functions of isolated proteins in non-physiologic models, and we lack scalable methods to extend domain shuffling to a broader set of proteins in primary human cells, where more complex functions and phenotypes could be explored^10–16^.

T cells have served as an exemplar primary human cell model for the development of functional genomics tools. CRISPR and related technologies have enabled genome-wide knock-out, knock-down, overexpression, and base-pair-specific tuning of natural genes, which has led to the discovery of basic biological mechanisms and nominated strategies for enhancing engineered T cell therapies^8,9,17–23^. As cardinal examples, overexpression of Activator Protein-1 (AP-1) transcription factors (TFs)—for example, *JUN*, *BATF*, *BATF3*, and *FOSL2*—can improve various T cell functions, including proliferation, cytokine production, stemness, or exhaustion resistance in the setting of chronic antigen stimulation^18,24–27^. While the DNA-binding domain (DBD) is largely conserved among AP-1 TFs, they differ in additional transactivation and protein interaction domains, which may be the result of gene duplication and subsequent divergence via domain-level changes^28–31^. The combinations of domains reflected in natural gene structures have proven utility in enhancing T cell function, but these arrangements are just a fraction of all possible combinations within the AP-1 family. Thus, non-evolved AP-1 domain configurations could unlock potentially latent functions that are not realized in current natural gene structures.

We developed a high-throughput combinatorial assembly method to generate thousands of barcoded novel genes from the domain building blocks of natural genes, termed DESynR (Domain Engineered via Synthesis and Recombination) genes. Within the AP-1 TF family, we generated 13,824 DESynR TFs from DNA-binding, transactivation, and protein interaction domains. The expression of DESynR AP-1 TFs in human chimeric antigen receptor (CAR) T cells significantly outperformed overexpression of their natural TF counterparts across various functions in *in vitro* and *in vivo* tumor challenge models. Patterns of domain utilization among top DESynR TFs was unexpected, including pairings across AP-1 subfamilies and with contributions from TFs that are minimally expressed in natural T cell subsets. Functional enhancements driven by DESynR TFs were linked to widespread transcriptional and epigenetic reprogramming, capable of inducing non-natural T cell states with gene expression programs from distinct non-T cell types. This approach introduces a new class of genetic perturbation that could be applied across protein families and cell types to comprehensively explore non-natural cell states and their therapeutic relevance.

### High-throughput assembly of novel AP-1 TF architectures via barcoded domain synthesis and recombination

To explore all possible domain combinations within a protein-coding gene family, we developed a method to iteratively assemble pools of domain sequences into a library of full-length, barcoded genes, termed DESynR (Domain Engineered by Synthesis and Recombination) genes (**Fig. 1a** and **Extended Data Fig. 1a**). By separating each domain sequence from a DNA-barcode (BC) with a restriction enzyme-excisable region, new domains could be added sequentially while also appending domain-associated BCs. Thus, after complete assembly, each gene could be identified by next-generation sequencing (NGS) of a short amplicon spanning the BCs. We first applied this method to AP-1 TFs, which exhibit a conserved basic leucine zipper (“bZIP”) DNA-binding and dimerization domain that is flanked to the “N” and “C” terminal ends by diverse additional domains, including transactivation and protein-interaction domains^28–31^. To encompass this diversity, we deconstructed 24 natural AP-1 TFs across the *ATF*, *FOS*, *JUN* and *MAF* subfamilies into three domains—“N”, “bZIP” and “C”—and synthesized 72 domains (ranging from 50-1,000 base pairs in length; **Fig. 1a,b**). Iterative combinatorial cloning of the “N”, “bZIP”, and “C” domain libraries created a full-length gene library of 13,824 DESynR AP-1 TFs (**Fig. 1c**). In contrast to individual gene synthesis, combinatorial construction decoupled library scale from synthesis costs and resulted in ∼100-fold reduction in cost.

**Figure 1.**
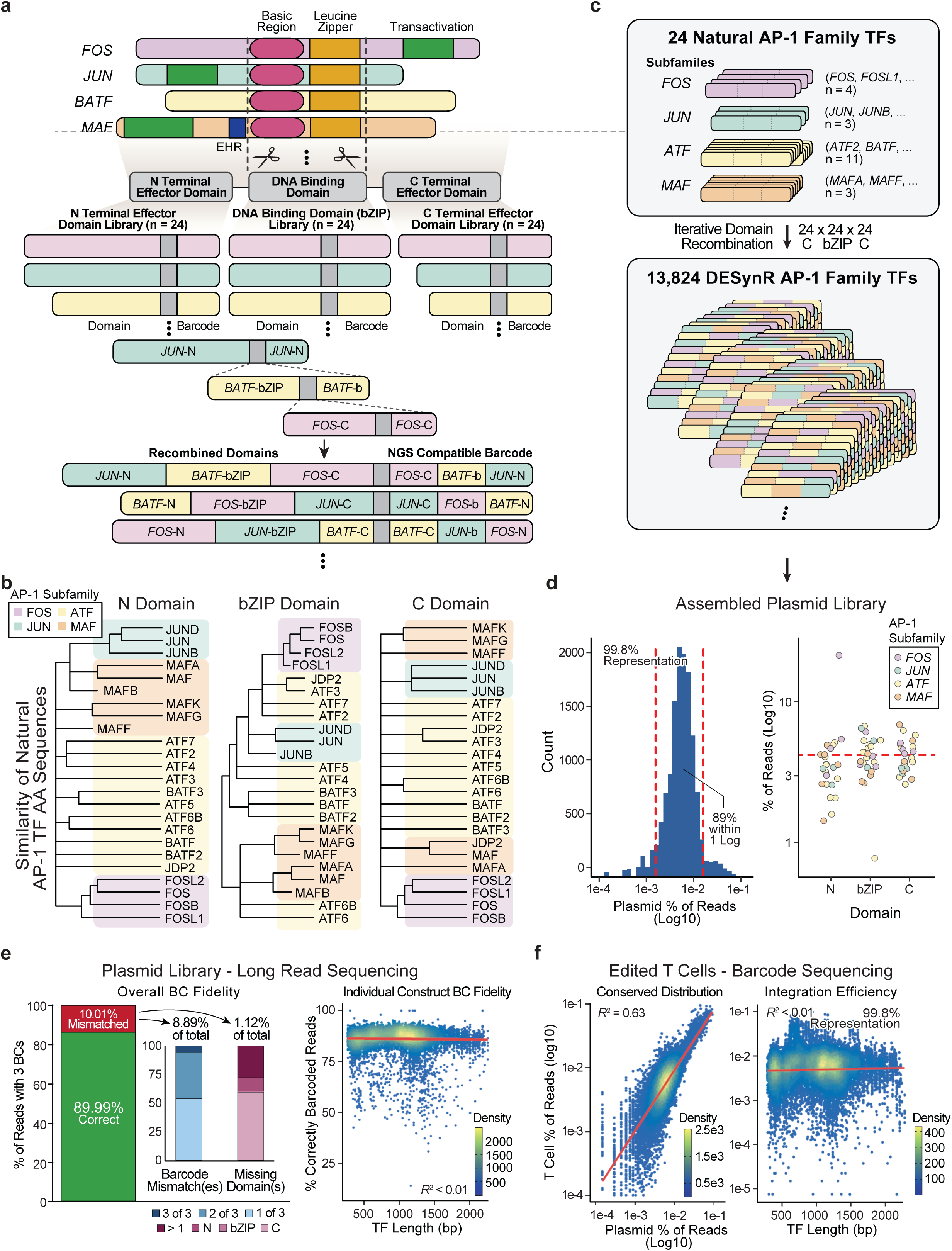
DESynR molecular platform for scalable, high-fidelity combinatorial assembly of new genes from constituent domains. **a)** Adapting characteristic AP-1 transcription factor domain structures (exemplar from each AP-1 subfamily shown) into barcoded domain pools for iterative combinatorial assembly into full-length DESynR TFs (EHR: Extended Homology Region)^45^. **b)** Alignment of natural AP-1 TF domain amino acid sequences by similarity using COBALT^46^. **c)** Applying the DESynR molecular platform to 24 natural AP-1 family TFs colored by subfamily generated 13,824 DESynR TFs reflecting diverse natural domain arrangements. **d)** Barcode (BC) sequencing of the assembled DESynR TF plasmid library. Left, a binned histogram to assess spread and representation. Right, a strip plot of each natural domain’s representation. **e)** Long-read sequencing of the final plasmid library, where each read spanned a single fully assembled DESynR AP-1 TF and its BC region to assess barcoding fidelity and cloning efficiency (filtered to reads with three BCs). Left, a stacked bar plot of overall (all constructs aggregated) TF-BC linkage fidelity and a breakdown of main error modes. Right, a scatter plot of the correlation in TF-BC linkage fidelity of individual constructs and base-pair (bp) length. Filtered to reads with three BCs and domains. R^2^ calculated from a y ∼ x linear regression. **f)** BC sequencing of edited T cells from a representative human donor (n=3). Left, a scatter plot to correlate BC representation in the final plasmid library versus edited T cells. Right, a scatter plot correlating TF representation to bp length. R^2^ calculated from a y ∼ x linear regression.

We first validated that the fully assembled DESynR AP-1 TF library achieved complete representation of domain combinations, high-fidelity BC linkages, and balanced constituent domain usage. Next-generation sequencing of the plasmid library showed 99.8% of all possible domain combinations were present and evenly distributed, as roughly 89% fell within one order of magnitude in their percentage representation (**Fig. 1d**). At the domain level, the 72 constituent domains were largely equally represented in the final library (**Fig. 1d**). To assess the fidelity of linkages between BCs and domains in assembled TFs, we performed long-read sequencing of four to six-kilobase (kb) amplicons from the plasmid library that included the fully assembled DESynR AP-1 TF and BCs. Of reads containing three valid BCs (∼95%), nearly 90% of the library was correctly linked (**Fig. 1e** and **Extended Data Fig. 1b,c**). The subset of mismatched reads (∼10%) included single, double, and triple domain-BC mismatches and instances of one or more missing domains (**Fig. 1e**). Importantly, barcoding errors did not correlate with TF size or lead to significant drop-out of specific DESynR TFs, as over 90% of the library maintained above 80% fidelity (**Fig. 1e** and **Extended Fig. 1d**). Finally, TF-BC linkages were orthogonally validated using whole-plasmid sequencing of 12 clones where 11/12 clones were correctly linked (**Extended Data Fig. 1e**).

Next, we validated that library quality was maintained upon genomic integration into primary human T cells. The DESynR AP-1 TF library was sub-cloned into a vector containing an FDA-approved CD19-28ζ CAR (axicabtagene ciloleucel; Yescarta) flanked by sequences with homology to the T-cell receptor α constant (*TRAC*) locus (**Extended Data Fig. 1a**). Upon electroporation of this plasmid library into T cells with a *TRAC*-cleaving CRISPR-Cas9 ribonucleoprotein, homology-directed repair resulted in an in-frame integration of the CAR and DESynR TF-containing multicistronic cassette into exon 1 of *TRAC*, which could then be transcribed using the endogenous *TRAC* promoter^32,33^.

Barcode sequencing of edited T cells showed that the distribution of the genomically integrated library was correlated with the plasmid pool (R^2^=0.63) and that representation of the DESynR TF library was high (>90%) in three human donors (**Fig. 1f** and **Extended Data Fig. 1f,g**). Additionally, the variable size of the DESynR AP-1 TFs did not impact integration efficiency (**Fig. 1f**). In sum, iterative and combinatorial construction of full-length genes from constituent domains generated a well-balanced and comprehensive DESynR library that retained barcoding fidelity and could be efficiently introduced into primary human T cells.

### DESynR AP-1 TFs outperform natural AP-1 TFs and enhance primary human T cell function

Next, we asked whether expression of DESynR TFs could drive distinct functional phenotypes in CAR T cells. Durable anti-tumor responses are limited by the intrinsic exhaustion of CAR T cells caused by chronic antigen stimulation, which results in diminished proliferation, persistence, effector function, and memory formation^34–38^. We specifically tested the ability of DESynR TFs to provide CAR T cells with a proliferative or persistence advantage in an *in vitro* exhaustion assay: a pool of DESynR AP-1 TF-harboring CAR T cells were repetitively challenged with CD19-expressing Nalm6 leukemia cells six times over the course of two weeks (**Extended Data Fig. 2a**). TF-barcode abundance changes—using amplicon sequencing of extracted genomic DNA— were assessed by comparing edited T cells from the input population to those at the end of: (1) the chronic stimulation assay, and (2) a parallel acute single stimulation assay to identify DESynR TFs that provided a general proliferative advantage, as well as exhaustion-specific enrichment. To technically validate the assays and BC sequencing workflow, we first screened a small subset of 216 DESynR AP-1 TFs generated via recombination of 3-9-member domain sub-libraries. We maintained high coverage of the library throughout cloning, T cell editing, and the exhaustion assay, and confirmed that all TF-BCs could be amplified from all three human donors (**Extended Data Fig. 2**).

Having validated the screening pipeline, we tested the comprehensive library of 13,824 DESynR AP-1 TFs along with 24 individually synthesized and barcoded full-length natural AP-1 TFs, three genes previously identified to improve T cell exhaustion as positive controls (*LTBR, IL2RA, TFAP4*) and “CAR Only” controls (n=4) without TFs (**Fig. 2a**)^18,23^. After chronic stimulation, 242 DESynR AP-1 TFs were significantly enriched (adjusted *p*<0.05), and surprisingly, 53 out-performed the top natural AP-1 TF, *JUNB* (*JUNB* Log_2_FC: 3.7; 53 DESynR TFs Log_2_FC range: 3.7-6.8; **Fig. 2b**). Enrichment of the top hits was consistent across three human donors, and there was a correlation (R^2^=0.58) between enrichment in acute and chronic screens, although a subset of hits was stimulation context-specific (**Fig. 2c** and **Extended Data Fig. 3**). Namely, relative to the top natural AP-1 TFs, 35 DESynR TFs were uniquely enriched in the chronic stimulation assay (Chronic Log_2_FC>3.7 [*JUNB*] & Acute Log_2_FC<2.8 [*BATF3*]) and 25 DESynR TFs were uniquely enriched in the acute stimulation assay (Acute Log_2_FC>2.8 [*BATF3*] & Chronic Log_2_FC<3.7 [*JUNB*]; **Fig. 2b** and **Extended Fig. 3c**). Top DESynR TFs comprised novel rearrangements of domains from natural AP-1 TFs previously identified as regulators of T cell exhaustion (*BATF3-FOSL1-JUN* Log_2_FC: 5.4, *JUN-ATF3-BATF3* Log_2_FC: 3.9), as well as rearrangements of domains from TFs previously undescribed in this context (*BATF2-JDP2-MAFK* Log_2_FC: 4.1, *FOSL1-ATF5-ATF6B* Log_2_FC: 3.8)^24,26^.

**Figure 2.**
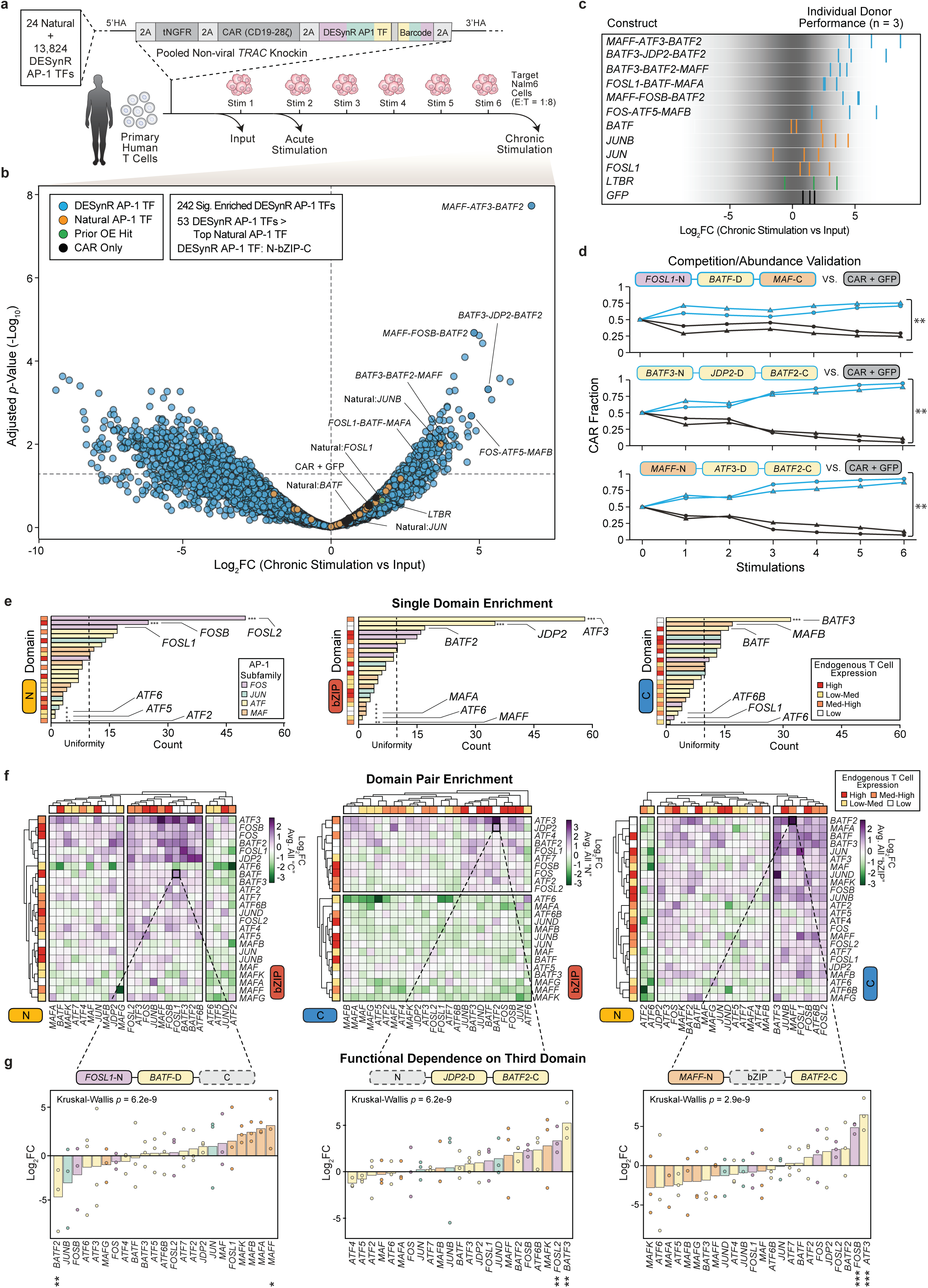
Pooled screening identifies DESynR TFs that outperform natural TFs and informs domain-use logic. **a)** DESynR TF pooled screening pipeline in primary human T cells. The library and controls were subcloned into a vector containing a CD19-28ζ CAR and a tNGFR reporter, flanked by homology to the *TRAC* locus to enable non-viral integration^32,33^. Barcodes from edited cells were sequenced before (input) and after a single stimulation (acute) or six stimulations over two weeks (chronic) with target Nalm6 leukemia cells at a 1:8 E:T ratio (see Methods). **b)** A volcano plot of the chronic stimulation screen in three human donors shows 242 significantly enriched DESynR TFs, including 53 DESynR TFs that outperformed the top natural AP-1 TF in the screen. Annotation of DESynR TFs: N-bZIP-C. Log_2_FC calculations were performed with DESeq2; significance was calculated by the Wald test and corrected for multiple comparisons using the Benjamini-Hochberg method. Hashed horizontal line indicates *p*=0.05. **c)** Log_2_FC in individual human donors for selected constructs. Each vertical bar represents one human donor (n=3). **d)** Three top performing DESynR AP-1 TFs (blue lines) from the pooled chronic stimulation screen were individually validated in a competitive chronic stimulation assay against CAR + GFP-control (black lines). Shapes indicate unique human donors (n=2). Independent two-tailed Student’s *t*-test performed after six stimulations. **e)** For 242 significantly enriched DESynR AP-1 TFs, bar plots enumerating utilization of “N”, “bZIP” and “C” domains from left to right. To the left of each bar plot is a heatbar of expression (colored by quartile) in endogenous T cells of the corresponding domain (see **Extended Data Fig. 4b**). Vertical dashed lines indicate uniform representation. Chi-squared tests were adjusted for multiple comparisons by FDR to determine significant deviations from uniformity. **f)** Each three-domain DESynR AP-1 TF was re-classified as three separate domain-pair architectures (“N+bZIP”, “bZIP+C” and “N+C”). Heatmaps for each architecture where each cell indicates the Log_2_FC of a unique domain-pair. Outside heatbars report expression of the domain’s corresponding full natural AP-1 TF in endogenous T cells (colored by quartile; see **Extended Data Fig. 4b**). **g)** Bar plots of Log_2_FC of exemplar domain-pairs completed with respective third domains. From left to right, “C” domain for “N+bZIP”, “N” for “C+bZIP,” and “bZIP” for “N+C”. Each dot represents one human donor (n=3). Log_2_FC calculations were performed with DESeq2; significance was calculated using the Wald test and corrected for multiple comparisons using the Benjamini-Hochberg method. Kruskal-Wallis (one-tailed) test used to assess overall deviation from uniformity. \**p*<0.05, \*\**p*<0.01, \*\*\**p*<0.0001.

**Figure 3.**
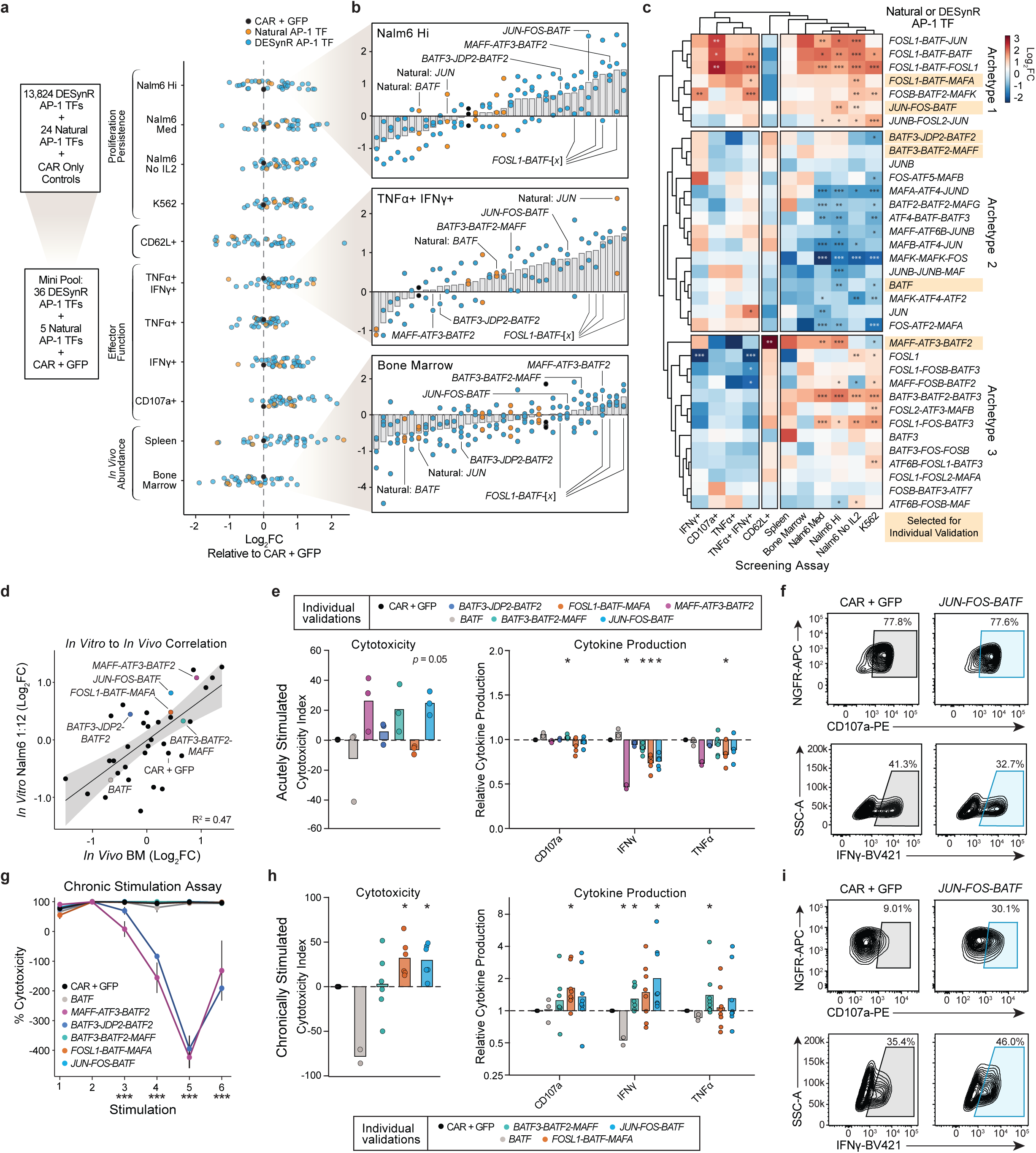
Multi-parameter pooled screening identifies functional DESynR TF archetypes. **a-c**, A mini-pool of CAR T cells edited with 36 DESynR AP-1 TFs, 5 natural AP-1 TFs and a GFP-control was screened across diverse *in vitro* and *in vivo* functional assays. Log_2_FC calculations were performed with DESeq2; significance was calculated by the Wald test and corrected for multiple comparisons using the Benjamini-Hochberg method (n=3 human donors for all “Proliferation/Persistence” and “CD62L+” screens, n=2 human donors for cytokine screens, and n=1 human donor and 3-4 mice for *in vivo* screens; see Methods). **a)** A strip plot of average Log_2_FC across donors scaled to CAR + GFP-control (except CD62L, see Methods). **b)** Bar plots of Log_2_FC from three highlighted functional screens with overlaid points representing one human donor/mouse. “*FOSL1*-*BATF*-[x]” indicates the “*FOSL1*-*BATF*” “N+bZIP” domain-pair with a variable “C” domain. **c)** Hierarchically clustered heatmap of average Log_2_FC. Highlighted TFs subjected to arrayed validation experiments. CAR + GFP-control omitted as not present in “CD62L” screen. **d)** Scatter plot of the correlation between Log_2_FC in *in vitro* “Nalm6 Hi” and *in vivo* “Bone Marrow” screens. R^2^ calculated from a y ∼ x linear regression. **e-i,** Arrayed validation experiments of highlighted DESynR AP-1 TFs, natural *BATF* and CAR + GFP-control. **e)** Left, a bar plot of *in vitro* acute cytotoxicity of Nalm6 target cells (CAR T cells not previously stimulated). Cytotoxicity index defined as cytotoxicity relative to CAR+GFP-control determined from a flow cytometric count of remaining Nalm6 target cells. Overlaid points represent average of technical duplicates or triplicates in three human donors (E:T=1:8). Right, a bar plot of degranulation and cytokine production following a single acute stimulation with Nalm6 target cells (CAR T cells not previously stimulated). Values are relative to CAR + GFP-control with overlaid points representing average of technical duplicates or triplicates across human donors (n=2-5). Significance was calculated by a paired two-tailed Student’s *t*-test. **f)** Exemplar flow cytometry plots of degranulation and IFNγ production from a representative human donor following a single acute stimulation (n=2-5 total donors). **g)** A line graph of CAR T cell cytotoxicity throughout an arrayed repetitive stimulation assay with six stimulations at a 1:8 E:T with Nalm6 target cells performed over two weeks. Each point represents mean +/-s.e.m. (n=2-4 human donors). One-way ANOVA performed for each stimulation to test for significant differences in cytotoxicity across constructs. **h)** Left, a bar plot of cytotoxicity index in a terminal challenge with Nalm6 target cells (E:T=1:12) following a two-week chronic stimulation assay. Overlaid points represent average of technical duplicates or triplicates (n=2-6 human donors). Right, a bar plot of degranulation and cytokine production in CAR T cells stimulated a final time following a two-week chronic stimulation assay. Values are relative to CAR + GFP-control with overlaid points representing average of technical duplicates or triplicates across human donors (n=2-8). Significance calculated by a paired two-tailed Student’s *t*-test. **i)** Exemplar flow cytometry plots of degranulation and IFNγ production from a representative human donor following chronic stimulation (n=2-8). \**p*<0.05, \*\**p*<0.01, \*\*\**p*<0.0001.

Select top-performers from the screen were individually fully synthesized (i.e. not constructed domain-by-domain) and sub-cloned into the CAR-vector for arrayed competitive chronic stimulation validation assays versus a CAR + GFP-control in two additional human donors (**Fig. 2d**). Mirroring their performance in the screen, proliferation of CAR T cell expressing each of these TFs was significantly greater than CAR + GFP-control T cells.

We then asked whether the enriched DESynR AP-1 TFs preferentially utilized particular single domains or pairs of domains (**Fig. 2e,f**). The *FOS* subfamily (*FOSL2*, *FOSB* and *FOSL1*) was significantly enriched (∼1.5-3x, *p*<0.05) at the “N” domain position (**Fig. 2e**). Domain usage at the “bZIP” position reflected amino acid sequence similarity; the top two significantly enriched domains (∼3-6x, *p*<0.0001), *ATF3* and *JDP2,* are highly similar, while significantly depleted domains (∼3-10x, *p*<0.05) across the *ATF* and *MAF* subfamilies also show sequence similarity (**Fig. 2e** and **Fig. 1b**). A dominant sub-family bias was not evident at the “C” domain position, although the structurally similar *BATF* and *BATF3* (∼3x enrichment, *p*<0.0001) were among the top three “C” domain hits (**Fig. 2e**). Strikingly, two (*JDP2, BATF2*) of the top three domains at the “bZIP” position and three (*BATF3, MAFB, MAFA*) of the top four domains at the “C” position came from TFs that are lowly expressed across endogenous T cell types, suggesting that the use of protein domains that are not typically expressed in T cells may drive novel cellular function (**Fig. 2e** and **Extended Data Fig. 4a,b**).

To assess abundance changes of domain-pairs in chronically stimulated versus input T cell populations, all DESynR TFs were re-classified as three separate domain-pairs (i.e. “N+bZIP,” “bZIP+C,” or “N+C”) for Log_2_FC calculations (**Fig. 2f** and **Extended Data Fig. 4c-f**). The majority of significantly enriched domain-pairs (48/67; adjusted *p*<0.05) featured at least one domain preferentially utilized by the 242 top three-domain performers, including *FOSL1-BATF* (N+bZIP), *ATF3-BATF* (bZIP+C), and *BATF3-BATF3* (N+C; **Extended Data Fig. 4d**). However, this was not a strict requisite as two domains, unenriched as single domains, could form a significantly enriched domain-pair (19/67; adjusted *p*<0.05), including *BATF3-FOSL1* (N+bZIP) and *MAFF-BATF2* (N+C; **Extended Data Fig. 4d**). Similar to single domain enrichment, across the three domain-pair architectures, the highest enrichment was observed in domain-pairs with one or both domains from lowly expressed AP-1 TFs, such as *BATF2-JDP2* (N+bZIP), *FOSL1-BATF3* (bZIP+C), and *BATF3-BATF2* (N+C; **Extended Data Fig. 4e**).

Finally, we asked whether all three domains were critical for functional outcomes. There was a very modest correlation (R^2^=0.07-0.12) between Log_2_FC of domain-pairs and full three-domain TFs; even highly enriched domain-pairs showed a broad range of enrichment as three-domain TFs depending on the identity of the third domain (**Extended Data Fig. 4f**). To confirm this global pattern, we closely examined a significantly enriched (*p*<0.05) exemplar domain-pair from each architecture (**Fig. 2g**). Indeed, enrichment was significantly impacted by the third domain although there were consistent patterns in the AP-1 sub-family identity of enriched or depleted third domains; for example, *MAF* family “C” domains were enriched in the *FOSL1-BATF* “N+bZIP” architecture (**Fig. 2g**). These results demonstrate that DESynR AP-1 TFs can enhance one aspect of T cell function— proliferation or persistence in the setting of chronic stimulation—beyond what is possible with natural AP-1 TFs. Notably, the top-performing DESynR TFs were not predictable based exclusively on the performance of their individual domains or domain-pairs, or their expression level or activity in endogenous T cells, underscoring the need for large-scale DESynR libraries.

### Multi-parameter functional assays uncover diverse DESynR AP-1 TF archetypes

We next assessed the impacts of DESynR AP-1 TFs on a broader set of T cell functions. We curated a list of 36 top-performing DESynR TFs from the exhaustion screen, including top TFs across diverse bZIP domains, and formed a mini-pool with five top natural AP-1 TFs and a CAR + GFP-control (**Fig. 3a**). For the mini-pool screens, all TFs were individually synthesized and assigned a new barcode before sub-cloning into the CAR-vector. Across a variety of functional assays *in vitro* and *in vivo*, we screened the mini-pool for: (1) proliferation and persistence in response to chronic antigen stimulation (*in vitro* repetitive stimulation assays & abundance in an *in vivo* Nalm6 leukemia xenograft model), (2) effector function (degranulation and IFNγ/TNFα production), and (3) retention of stemness (CD62L expression; **Extended Data Fig. 5a**). Analysis of *in vitro* chronic stimulation assays confirmed results from the prior screens, as the majority (>50%) of DESynR TFs out-performed natural AP-1 TFs and CAR + GFP-control when chronically exposed to Nalm6 cells (**Fig. 3a,b**). Top performers were largely consistent across proliferation and persistence assays, although there was some variability in variations of the assay (i.e. absence of exogenous IL-2 and stimulation with a different leukemic cell line), underscoring the subtle influences of environment on the proliferative phenotype. For example, the top hit in the large screen, *MAFF*-*ATF3*-*BATF2*, as well as the *FOSL1*-*BATF* “N+bZIP” domain-pair, showed increased proliferation across assays. Across effector function screens, the majority of DESynR TFs (∼50-90% of constructs), particularly those containing the *FOSL1*-*BATF* “N+bZIP” domain-pair, out-performed the CAR + GFP-control, while natural AP-1 TFs performed variably (natural AP-1 TFs made up ∼12% of the mini-pool but <5% of the top quartile of effector function hits). *MAFF*-*ATF3*-*BATF2* was the consensus winner of the stemness screen (Log_2_FC=2.4, *p*<0.001). Finally, *in vivo* screens in a Nalm6 leukemia xenograft model showed enrichment of both natural and DESynR AP-1 TFs in the spleen ten days after adoptive transfer, with a smaller set (n=11) of exclusively DESynR AP-1 TFs, in particular *FOSL1*-*BATF* “N+bZIP” domain-pairs, enriched in the bone marrow (**Fig. 3b** and **Extended Data Fig. 5b-d**).

To understand whether groups of DESynR AP-1 TFs induced similar patterns of T cell functional behavior, we constructed a genotype-phenotype map of natural and DESynR AP-1 TFs across the array of functional screens (**Fig. 3c**). Notably, we identified three distinct archetypes of TFs based on their functional performance. TFs with the highest effector function and proliferative capacity and relatively decreased stemness formed Archetype 1. The more diverse Archetype 2 TFs were largely poor proliferators with variable effector function and stemness. Finally, Archetype 3 TFs retained stemness without loss in proliferative capacity. Strikingly, Archetype 1 was exclusively occupied by DESynR AP-1 TFs, suggesting that DESynR TFs could establish a novel T cell phenotype that more readily accesses a program of key effector functions (cytokine production and proliferation) than natural AP-1 TFs.

Next, we individually validated the phenotypes observed in the mini-pool screens and tested cytotoxicity, which is a requisite function for therapeutic T cells, but difficult to measure via pooled screening. We selected at least one DESynR AP-1 TF from each Archetype (highlighted in **Fig. 3c**) that was also enriched in the bone marrow *in vivo* (annotated in **Fig. 3d**). Since each of these candidates featured a domain from *BATF* or the greater *ATF* subfamily, and *BATF* overexpression has previously been demonstrated to improve CAR T cell function, we chose *BATF* as a control natural AP-1 TF^18,25^. We assessed cytotoxicity following acute stimulation, throughout repetitive rounds of stimulation, and in a terminal challenge following repetitive stimulation. Acutely, across a range of effector to target ratios (E:Ts), CAR T cells expressing DESynR TFs killed Nalm6 tumor cells as well as or better than CAR + GFP-control cells or *BATF*-overexpressing CAR T cells, despite generally secreting slightly less effector cytokines (**Fig. 3e** and **Extended Data Fig. 6**). Expression of several inhibitory receptors (IRs -PD1, LAG3, CD39) was largely unchanged in CAR T cells with DESynR TFs, and *FOSL1*-*BATF*-*MAFA*-edited cells expressed higher (∼1.5x) levels of stemness markers, CD45RA and CD62L (**Extended Data Fig. 6d,g**). Unexpectedly, by the third stimulation of the chronic stimulation assay, there was significant (*p*<0.0001) divergence in cytotoxicity, and CAR T cells expressing *MAFF*-*ATF3*-*BATF2* and *BATF3*-*JDP2*-*BATF2* lost control of Nalm6 targets, while CAR T cells expressing *FOSL1-BATF-MAFA*, *BATF3-BATF2-MAFF*, *JUN-FOS-BATF*, and those overexpressing natural *BATF* consistently killed all Nalm6 targets; this cytotoxicity defect was not accompanied by reduced proliferation (**Fig. 3g** and **Extended Data Fig. 6a**).

After two weeks of chronic stimulation, we challenged T cells with a terminal cytotoxicity assay across a range of E:T ratios. Here, CAR T cells expressing two DESynR AP-1 TFs, *FOSL1*-*BATF*-*MAFA* and *JUN*-*FOS*-*BATF* (both *p*<0.05), consistently out-performed CAR + GFP-control and overexpression of the natural AP-1 TF *BATF* (**Fig. 3h** and **Extended Data Fig 6i**). In parallel, DESynR AP-1 TFs showed significantly (*p*<0.05) increased degranulation (*FOSL1*-*BATF*-*MAFA*) and effector cytokine secretion (IFNγ - *JUN-FOS-BATF*, *BATF3-BATF2-MAFF*; TNFα - *BATF3-BATF2-MAFF*) following chronic stimulation, mirroring results from the mini-pool screens (**Fig. 3h**). CAR T cells expressing DESynR TFs generally downregulated IRs relative to CAR + GFP-control, while cells expressing *FOSL1*-*BATF*-*MAFA*, consistently showed an increased frequency of central-memory (CD45RA-CD62L+) T cells (**Extended Fig. 6e,f**).

Finally, we tested CAR T cells expressing the top performing DESynR AP-1 TFs against overexpression of the natural AP-1 TF *BATF* in an *in vivo* Nalm6 xenograft leukemia model. Similar to their improved performance in the terminal cytotoxicity and cytokine production assays, CAR T cells expressing *JUN*-*FOS*-*BATF* and *FOSL1*-*BATF*-*MAFA* significantly extended overall survival compared to CAR + GFP-control T cells (*p*<0.05) or overexpression of the natural AP-1 TF *BATF* (**Extended Data Fig. 7**). Overall, deeper phenotyping of DESynR AP-1 TF-edited CAR T cells uncovered differential engagement of cardinal T cell functions and nominated a functional Archetype (1) exclusive to DESynR TFs that may hold therapeutic promise.

### DESynR TFs program non-natural transcriptional and epigenetic T cell states

Since DESynR AP-1 TFs could enhance CAR T cells across a broad range of functions, we asked whether these phenotypes were linked to underlying large-scale epigenetic and transcriptional changes. First, we performed single-cell RNA sequencing (scRNA-seq) of *in vitro* chronically-stimulated CAR T cells expressing either a DESynR AP-1 TF (*BATF3*-*JDP2*-*BATF2, MAFF*-*ATF3*-*BATF2, JUN*-*FOS*-*BATF, BATF3-BATF2-MAFF*, *or FOSL1-BATF-MAFA*), a natural AP-1 TF (*BATF* or *JUNB*), or CAR + GFP-control (n=5 human donors; **Fig. 4a** and **Extended Data Fig. 8)**. Unsupervised clustering of cells (n=52,952) integrated by human donor generally segregated cells into clusters marked by *CD4* or *CD8A*, and proliferative (*MKI67*, *HIST1H3G*) or cytotoxic (*PRF1, GZMB*) phenotypes, along with a Treg-like cluster (*FOXP3*, *CTLA4*; **Fig. 4a,b** and **Extended Data Fig. 8k**). Notably, several clusters were almost exclusively comprised of cells expressing DESynR TFs (**Fig. 4c,d** and **Extended Data Fig. 8l**). Specifically, nearly all *MAFF*-*ATF3*-*BATF2* and *BATF3*-*JDP2*-*BATF2* cells occupied proliferative and cytotoxic CD4+ clusters that were depleted of CAR + GFP-control and natural AP-1 TF cells (**Fig. 4d** and **Extended Data Fig. 8l**). Roughly half of *JUN*-*FOS*-*BATF*-expressing cells also occupied these DESynR TF-specific clusters, although *JUN*-*FOS*-*BATF*-expressing cells were also represented in a broader array of clusters overlapping with CAR + GFP-control and natural AP-1 TF-expressing cells (**Fig. 4d** and **Extended Data Fig. 8l**). Moreover, these three DESynR TFs were relatively depleted from the Treg-like cell cluster (**Fig. 4d**). Overall, *MAFF*-*ATF3*-*BATF2* (4,207 differentially-expressed genes, DEGs), *BATF3*-*JDP2*-*BATF2* (4,667 DEGs), and *JUN*-*FOS*-*BATF* (2,460 DEGs) drove more substantial transcriptional changes relative to CAR + GFP-control than either natural AP-1 TF, *JUNB* (1,045 DEGs) or *BATF* (1,537 DEGs), while *BATF3-BATF2-MAFF* (1,269 DEGs) and *FOSL1-BATF-MAFA* (1,169 DEGs) were roughly consistent with the natural AP-1 TFs (DEGs: absolute Log_2_FC>0.5 & exp. in >5% cells; **Extended Data Fig. 8g**). The extent of transcriptional remodeling was not tightly linked to functional Archetype identity, and functional enhancement could be provided by both TFs that were transcriptionally more similar to (*FOSL1-BATF-MAFA*) and divergent from (*JUN*-*FOS*-*BATF*) CAR + GFP-control (**Fig. 3c**).

**Figure 4.**
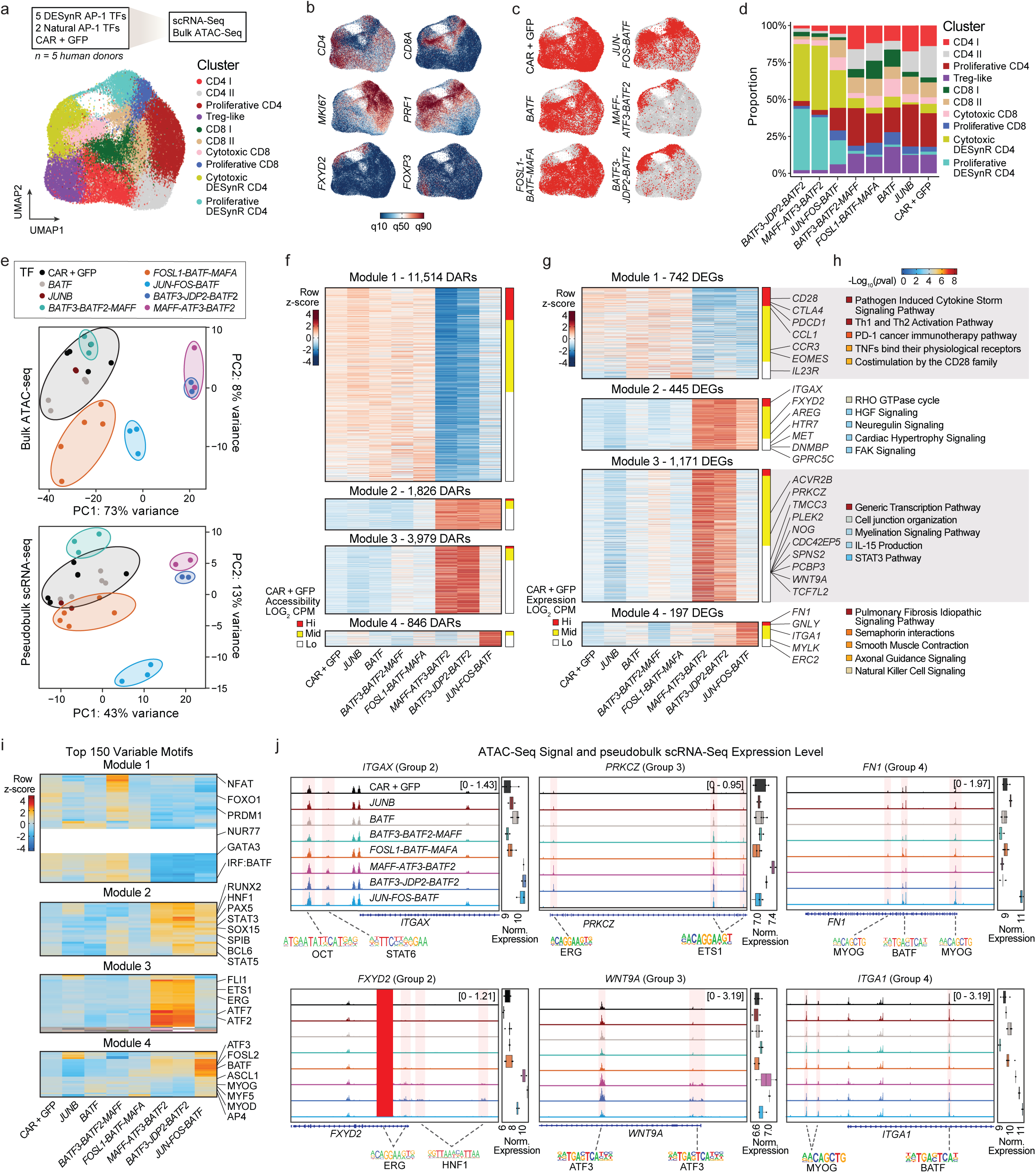
DESynR TFs program novel T cell states. **a-d**, Single-cell (sc) RNA-seq of 5 DESynR AP-1 TFs, 2 natural AP-1 TFs, and CAR + GFP-control CAR T cells following two weeks of arrayed chronic stimulation with Nalm6 leukemia target cells (n=3-5 human donors per construct; 52,952 total cells). **a)** Uniform manifold approximation and projection (UMAP) of donor-integrated data across all constructs. Ten clusters were identified by the Louvain method and annotated manually. **b)** Scaled expression of selected marker genes. **c)** UMAPs with red dots indicating cells of selected constructs with all other cells as gray dots, highlighting DESynR AP-1 TF-specific regions of the UMAP. **d)** A stacked bar plot showing occupancy in each annotated cluster for all constructs. **e-j,** Pseudobulk RNA-seq data and bulk ATAC-seq data following arrayed chronic stimulation (n=3-5 human donors per construct). **e)** Top, principal-component analysis (PCA) of the top 2,000 most variable chromatin regions from ATAC-seq data. Bottom, PCA of the top 1,000 most variable genes from pseudobulk RNA-seq data. **f)** Hierarchically clustered (k-means) heatmap of 18,165 differentially accessible peaks (DARs) relative to CAR + GFP-control from bulk ATAC-seq data. Top color bar indicates normalized z-scores for each DAR. Each module is ordered by accessibility (high to low) in CAR + GFP-control as shown by the bottom heatbar; all DARs are considered and heatbar color indicates top, middle and bottom third of accessibility. **g)** Hierarchically clustered (k-means) heatmap of differentially expressed genes (DEGs) relative to CAR + GFP-control from pseudobulk RNA-seq data. Top color bar indicates normalized z-scores for each DEG. Each module is ordered by expression (high to low) in CAR + GFP-control as shown by the bottom heatbar; all DEGs are considered and heatbar color indicates top, middle and bottom third of expression. **h)** Curated upregulated pathways determined by QIAGEN Ingenuity Pathway Analysis (IPA). Heatbar of -Log_10_ adjusted *p*-value. **i)** Heatmap showing chromVAR bias-corrected ATAC-seq deviations of the top 150 most variable TF motifs across all constructs, clustered using k-means unsupervised analysis. Color bar indicates normalized z-scores for each motif. **j)** Chromatin accessibility tracks for selected significantly different loci along with normalized gene expression (pseudobulk RNA profiles). Average of human donors (n=3-5) for both data types.

Next, we performed ATAC-seq on the same cell populations to assess global changes in chromatin accessibility (**Fig. 4e** and **Extended Data Fig. 9**). Similar to transcriptional changes, DESynR TFs, *MAFF*-*ATF3*-*BATF2* (15,317 differentially accessible regions, DARs), *BATF3*-*JDP2*-*BATF2* (14,129 DARs)*, JUN*-*FOS*-*BATF* (3,235 DARs), and *FOSL1-BATF-MAFA* (1,332 DAR) drove more substantial epigenetic changes relative to CAR + GFP-control than the natural AP-1 TF, *BATF* (679 DARs) or *BATF3-BATF2-MAFF* (182 DARs; DARs: absolute Log_2_FC>0.5 & FDR<0.05; **Extended Data Fig. 9d**). Principal Component Analysis (PCA) of the top 2,000 most variable chromatin regions showed that the strongest variance (PC1:73%) was between cells expressing CAR + GFP-control or natural *BATF* or *JUNB* (which clustered together), and cells expressing *MAFF*-*ATF3*-*BATF2* or *BATF3*-*JDP2*-*BATF2* (**Fig. 4e**). PC1 also partially segregated cells expressing *JUN*-*FOS*-*BATF*, which additionally varied along PC2 (8%), as did *FOSL1-BATF-MAFA* (**Fig. 4e**). For direct comparisons to transcriptional data, we generated pseudobulk profiles from the scRNA-seq data, and PCA of the top 1,000 most variable genes showed a very similar pattern: a dominant variance (PC1:43%) of *MAFF*-*ATF3*-*BATF2* and *BATF3*-*JDP2*-*BATF2* that was partially present in *JUN*-*FOS*-*BATF*, and a unique variance of *JUN*-*FOS*-*BATF* along a less dominant (11%) PC (**Fig. 4e**). Pseudobulk RNA profiles from CD4 and CD8 clusters confirmed that the distinct transcriptional states were driven by the DESynR TF and consistently observed in both CD4+ and CD8+ cells (**Extended Data Fig. 9a,b**).

We then defined variable chromatin regions relative to CAR + GFP-control and identified four modules of DARs (**Fig. 4f**). Similar analysis of pseudobulk RNA profiles yielded an analogous pattern of modules of differentially expressed genes (**Fig 4g**). We performed a permutation test to determine whether DARs were significantly enriched near genes in the corresponding gene expression module compared to a random background gene set and found significant enrichment for each module (*p*<0.0001; **Extended Data Fig. 9f**). Module 1, specific to CAR + GFP-control cells and cells expressing natural AP-1 TFs (*BATF* or *JUNB*) or two of the DESynR TFs (*BATF3-BATF2-MAFF* or *FOSL1-BATF-MAFA*), was characterized by genes and chromatin regions commonly associated with antitumor T cell responses and exhaustion (*CTLA4*, *PDCD1*, *EOMES*, *BTLA*)^37,39–41^. This module showed lower gene expression and accessibility in cells expressing the other three DESynR TFs (*MAFF*-*ATF3*-*BATF2*, *BATF3*-*JDP2*-*BATF2*, or *JUN*-*FOS*-*BATF*).

Conversely, the other modules (modules 2-4) were exclusive to *MAFF*-*ATF3*-*BATF2*, *BATF3*-*JDP2*-*BATF2*, and *JUN*-*FOS*-*BATF*-expressing cells. Some of the genes in modules 2-4 are common across all cell types or specific to T cell function; for example, module 4, exclusive to *JUN*-*FOS*-*BATF*, was enriched in cytotoxicity genes (*PRF1, GNLY, GZMB, GZMH*), which aligns with the functional enhancement that the DESynR TF provides.

Unexpectedly, we noted that many of the genes in modules 2-4 are typically expressed in disparate non-immune cell tissues (**Extended Fig. 9f**). To understand whether the altered chromatin accessibility and gene expression driven by DESynR AP-1 TFs in modules 2-4 acted on existing T cell programs or increased access to and transcription of new programs not typically expressed in CAR T cells, we rank-ordered all DARs and DEGs by absolute accessibility and expression, respectively, in CAR + GFP-control cells (**Fig. 4f,g**). Strikingly, DARs in modules 2-4 were predominantly lowly accessible in CAR + GFP-control cells, indicating that DESynR TFs can increase accessibility of chromatin regions that are naturally lowly or not accessible (**Fig. 4f**). Analysis of DEGs yielded a similar result: 1.5-2.5x more genes were lowly expressed by CAR + GFP-control cells in the DESynR TF-specific modules (**Fig. 4g**). We then performed biological pathway analysis to identify signatures of newly expressed gene programs. Module 2 was enriched in pathways such as “Neuregulin Signaling” and “Cardiac Hypertrophy Signaling”, characterized by accessibility and expression of non-T cell genes, including *AREG, ITGAX* and *PRKD3*, and enrichment of motifs of PAX5 and BCL6 (**Fig. 4f-j**). Module 3 was associated with “Myelination Signaling Pathway”, driven by increased accessibility and expression of *ACVR2B*, *TCF7L2* and *WNT9A* and enrichment of ETS-family motifs (**Fig. 4f-j**). Additionally, module 3 was characterized by T cell memory-associated processes such as “STAT3 Signaling” and “IL-15 Signaling”^42,43^. Finally, module 4 was enriched in pathways including “Smooth Muscle Contraction” and “Axonal Guidance Signaling”, marked by accessibility and expression of *GUCY1A1*, *ITGA1* and *MYLK* and enrichment of motifs such as MYOG, MYOD and ASCL1, typically associated with neuronal and muscular processes (**Fig. 4f-j**). Taken together, DESynR AP-1 TFs can induce epigenetic and transcriptional remodeling that program T cells to dampen the effects of T cell exhaustion, but more broadly, access non-natural cell states characterized by chromatin accessibility and gene expression from disparate non-immune tissues.

## Discussion

The DESynR platform enables the modular assembly and screening of thousands of novel genes constructed via the rearrangement of existing domains, mimicking evolutionary gene creation principles outside of organism-level selection constraints, and thereby enabling the identification of the ideal genes for a specific cellular function. This platform innovates in two significant ways: (1) overcomes the prohibitive costs of assembly of large combinatorial domain libraries using standard DNA synthesis, and (2) introduces a facile and flexible genetic barcoding strategy for screening newly constructed genes in any primary cell context. The latter may be particularly useful in assessing the impacts of DESynR genes with single-cell genomics to develop genotype-phenotype maps. Although we focused here on the generation of DESynR TFs within the AP-1 family, we envision that this technology could be seamlessly applied to build new genes from natural domains within other families, across multiple families, and in combination with synthetic domains^44^.

Since domain recombination can traverse far greater sequence design space than single base-pair mutations or short insertions or deletions, we hypothesized that DESynR TFs may be able to program non-natural cell states with distinct functional phenotypes^11^. This may be particularly relevant for settings where natural cell states have not achieved an optimal solution for a particular function given opposing selection pressures; for example, T cell exhaustion evolved to balance effector function and auto-inflammation in the setting of chronic antigen^37–40^. Indeed, in the models used here, DESynR AP-1 TFs outperformed their natural counterparts across diverse functional axes *in vitro* and *in vivo* and were able to program a distinct T cell functional archetype that could maintain persistence and effector function in the setting of exhaustion. In some cases, this was linked to broad epigenetic and transcriptional remodeling that established non-natural T cell states marked by gene expression programs borrowed from non-immune tissues. Future work will leverage the DESynR platform to build an expansive library of non-evolved genes, which can be screened in diverse cell types and biological contexts to create an atlas of non-natural cell states that can inform both basic biology as well as therapeutic applications.

## Supplementary tables

S1. DNA Sequences

S2. Pilot screen NGS read counts & results generated with DESeq2.

S3. Full screen NGS read counts & results generated with DESeq2.

S4. Mini-pool screens NGS read counts & results generated with DESeq2. S5. Antibodies

## Methods

### DNA Constructs and HDR Templates

#### Pooled Cloning

Each individual barcoded TF domain or full-length TF sequence (native AP-1 control sequences, **Supplementary Table 1**) was codon optimized, commercially synthesized (Twist Bioscience) and cloned into a Kanamycin-resistant vector (pTwist Kan vector). Individual plasmids were combined into equimolar pools of “N”, “bZIP” and “C” domain sequences or of full-length TF sequences. The domain-by-domain full-length gene library was built using iterative rounds of restriction digestions, ligations and bacterial transformation (Fig 1a). 1 ug of plasmid pools containing TF domains were digested in two steps: first with 2.5 uL of BsaI restriction enzyme (New England Biolabs, NEB) in 50 uL total volume (with CutSmart Buffer, NEB) for 1 hour at 37°C and then 2.5uL of SrfI was added and incubation was continued for a second hour. The destination backbone plasmids were digested with BbsI and PmeI (NEB) using the same protocol. Digestion products were SPRI-purified at a 2X volume ratio and eluted in H_2_O. Digested domain inserts were ligated into digested destination plasmid backbones using T7 ligase and StickTogether DNA Ligase Buffer (both NEB) at a 1:1 molar ratio with 20 ng of digested backbone per 200 uL ligation reaction for 30 minutes at room temperature (RT). Ligation products were SPRI-purified at a 1X volume ratio and eluted in H_2_O. Up to 100 ng of purified ligation product was electroporated into Endura electrocompetent cells (Lucigen) using a Gene Pulser XCell instrument (Biorad), recovered for 1 hour at 30°C with shaking and then grown for 12-16 hours at 30°C prior to plasmid preparation using ZymoPURE II kits (Zymo Research). To generate the final homology-directed repair template (HDRT) library, the full-gene library containing assembled N, bZIP, and C domain pools was similarly cloned into a vector containing a CD19-28ζ CAR and sequence homology to the *TRAC* locus (CAR vector, **Supplementary Table 5**). The HDRT library was eluted in H_2_O and endotoxin removal (Zymo) was performed. For the plasmid library of synthesized full-length TF sequences (i.e. not individual domains), a single digestion and ligation into the CAR vector, bacterial transformation and plasmid preparation was performed by the outlined methods to generate the HDRT plasmid library. The sequences of a sample of individual clones from both libraries were confirmed by whole plasmid next-generation sequencing (Plasmidsaurus).

#### Arrayed Cloning

For validation and mini-pool screening, full-length TF sequences (**Supplementary Table 1**) were individually commercially synthesized as Gene Fragments (Twist Bioscience). Using the same digestion and ligation protocols as for “*Pooled Cloning*”, these sequences were cloned into the CAR vector to generate HDRTs. Purified ligations were transformed into chemically competent cells (Mix & Go! Competent Cells - DH5 Alpha, Zymo Research). Sequences were confirmed by whole plasmid next-generation sequencing (Plasmidsaurus) before large-scale plasmid preparations were made using ZymoPURE II Kits (Zymo Research). The plasmids were eluted in H_2_O and endotoxin removal (Zymo) was performed.

### Primary Human T Cell Isolation, Activation and Culture

Leukoreduction chambers from processing platelet donations, collected under an approved IRB protocol from healthy human donors, were acquired from the Stanford Blood Center. Per manufacturer instructions, PBMCs were isolated by density centrifugation (Lymphoprep, StemCell) and subsequently CD3+ T cells negatively selected using the Human CD3 T Cell Enrichment Kit (StemCell). Following an automated count (Countess, Thermo Fisher) CD3 positive T cells were activated using CD3/28 Dynabeads (Thermo Fisher) at a one T cell per bead ratio and 1e6 T cells/mL in T cell medium (TCM): X-VIVO 15 medium (Lonza) supplemented with 5% FBS and 50 IU/mL human IL-2 (Peprotech). For *in vivo* experiments, CD3+ T cells were negatively selected from leukapheresis packs (StemCell) that were thawed after previous cryopreservation. Isolated T cells were counted and activated as outlined above, but in a distinct medium: X-VIVO 15 medium (Lonza) supplemented with 5% human serum (Gemini), 1% Penicillin-Streptomycin (Gibco) and 5 ng/mL IL-7 and IL-15 (Miltenyi).

### T Cell Editing

48 hours following activation, Dynabeads were magnetically removed (EasySep Magnet, StemCell) and T cells counted. T cells were centrifuged for 10 minutes at 90g and re-suspended in 1-2e6 T cells per 20 uL P3 Buffer (Lonza) and mixed with prepared Cas9 ribonucleoprotein (RNP) and HDRT. RNP was prepared such that for each editing reaction, 0.375 uL of 200 uM tracrRNA (Integrated DNA Technologies, IDT) was first mixed with 0.375 uL of 200 uM crRNA (IDT) and incubated for 15 minutes at RT to complex the gRNA. Then, 0.25 uL of 100 mg/mL 15-50 kDa poly-L-glutamic acid (Millipore Sigma) was added to the gRNA and mixed. Finally, 0.5 uL of 40 uM spCas9 (UC Berkeley MacroLab) was mixed in and incubated for 15 minutes at RT. The complexed RNP, 2 ul of 1 ug/uL plasmid HDRT and 20 uL of T cell suspension were mixed and aliquoted into a 96-well Nucleocuvette Plate (Lonza). Electroporation occurred on a Gen2 Lonza 4D instrument with a 96-well plate attachment using the pulse code EO-151. Immediately after electroporation, 80 uL of pre-warmed TCM was gently added to each well and incubated for 15 minutes at 37°C. The cells were then transferred to standard 96-well round-bottom plates with 300 uL total TCM. T cells were maintained at 0.5e6-1e6/mL and TCM was refreshed every two to three days. T cell editing for *in vivo* experiments was performed identically, except for the T cell medium used (see **Primary Human T Cell Isolation, Activation and Culture**).

The 20 bp *TRAC* exon targeting sequence of the Cas9 crRNA used in all experiments: 5’ - AGAGTCTCTCAGCTGGTACA - 3’.

### Cell Lines

Nalm6 cells were obtained from American Type Culture Collection (ATCC) and maintained in RPMI (Fisher Scientific) supplemented with 10% FBS. A distinct set of Nalm6 cells were previously engineered to express firefly luciferase-GFP (FFLuc-GFP)^49^. These cells were maintained in RPMI (Thermo Fisher) supplemented with 10% and 1% penicillin-streptomycin (100 U/mL, Thermo Fisher). K562 engineered to express human CD19 were a kind gift from the Roybal Lab (UCSF) and maintained in RPMI + 10% FBS.

### Flow Cytometry and Cell Sorting

Unless otherwise noted, cells were stained in 96-well round-bottom plates at RT in 20 ul total volume for 20”. Antibody (**Supplementary Table 5**) staining mix was made in FACS buffer: 1x phosphate-buffered saline (PBS) without calcium and magnesium supplemented with 2% FBS and 1 mM EDTA. FACS buffer was also used for washing cells post-staining and resuspension pre-acquisition. Stained cells were acquired on a Biorad ZE5 Cell Analyzer or BD LSRFortessa X-50 and analysis was performed using FlowJo software (BD). Cell sorting was performed using a FACSAria II (BD) equipped with a 70-um nozzle at RT. Cells were sorted into X-VIVO 15 medium supplemented with 50% FBS.

### In Vitro Assays

#### Repetitive Stimulation Assay

Starting on day 6 post T cell editing (Day 8 post initial T cell activation), CAR T cells were co-cultured in TCM with Nalm6 cells at effector to target (E:T) ratios indicated in figure legends at a density of ∼0.5e6 cells/cm^2^ and ∼1e6 cells/ml (**Extended Data Fig. 2a**). Every two to three days (Day 10, 13, etc. post initial T cell activation), the co-culture(s) was sampled (50 uL in duplicate/triplicate) and stained with antibodies against NGFR (CAR) and CD19 (Nalm6) and a viability dye (**Supplementary Table 5**). Samples were acquired with CountBright Plus Absolute Counting Beads (Thermo Fisher) to determine the counts of CAR T cells and Nalm6 cells. The co-culture was re-seeded to the predefined E:T.

#### Competition Assay

The “*Repetitive Stimulation Assay*” was performed with a 1:1 mix of T cell populations: one population edited with a TF construct (prepared according to “*Arrayed Cloning*”) and the second edited with GFP (CAR + GFP control). When assessing the co-cultures by flow cytometry for re-seeding, the CAR (NGFR+) population was sub-gated based on GFP status to determine the proportion of each T cell population.

#### Cytokine Assays

Arrayed cytokine assays were performed at baseline (Day 6, prior to “*Repetitive Stimulation Assay*”) or following the assay (Day 23, **Extended Data Fig. 2a**). CAR T cells were co-cultured with Nalm6 cells at 1:1-1:2 E:T ratios in 96-well round-bottom plates in 200 ul TCM without IL-2, supplemented with GolgiStop and GolgiPlug (Becton Dickinson - BD) for four to six hours. Intracellular staining was performed using the Cytofix/Cytoperm Kit (BD) using antibodies against IFNγ and TNFα following standard surface staining (“*Flow Cytometry”*) of CD4, CD8, NGFR and CD19 along with a viability dye (**Supplementary Table 5**). Stained cells were acquired on a Ze5 Cell Analyzer (Biorad) and analysis was performed using FlowJo software (BD).

#### Cytotoxicity Assays

Arrayed cytokine assays were performed at baseline (Day 6, prior to “*Repetitive Stimulation Assay*”) or following the assay (Day 23, **Extended Data Fig. 2a**). 10,000 CAR T cells were seeded in 96-well round-bottom plates with variable numbers of Nalm6 cells as indicated in figure legends in 200 ul total TCM in duplicate or triplicate. After 24 hours, co-cultures were analyzed by flow cytometry using counting beads as described in “*Repetitive Stimulation Assay”*.

### T Cell Screening

For pooled screening using the “*Repetitive Stimulation Assay”*, T cells were expanded for six days post-electroporation. gDNA was extracted from >500 edited T cells/construct prior to the start of the assay (Input), following one stimulation (Acute) and after six stimulations (Chronic) using NucleoSpin Blood kits (Macherey-Nagel) (**Extended Data Fig. 2a**). Several variations of the “*Repetitive Stimulation Assay”* were used for mini-pool screening (Fig. 3 and **Extended Data Fig. 5a**) including different E:Ts, without IL-2 supplementation and with a different target cell line (K562-CD19 1:1 in TCM).

At the end of the Nalm6 1:10 screen, chronically stimulated cells were also stained for CD62L expression and cytokine production (**Extended Data Fig. 5a**). For CD62L, cells were stained with an anti-human CD62L antibody, the population was sorted into the top

∼25% and bottom ∼25% of CD62L expression and gDNA was extracted. For cytokine, cells were stimulated and stained (“*Cytokine Assays*”) and four-way sorted (IFNγ+, TNFα+, double-positive and double-negative) and gDNA was extracted with a modified protocol. Briefly, cell pellets were resuspended in 200 uL diH_2_O plus 4 uL 0.5 M pH 8 EDTA, 8 uL 1M pH 8 TRIS and 2 uL 20 mg/mL Proteinase K (Thermo Fisher) and incubated for 12-16 hours at 55°C with agitation. Then, genomic DNA was extracted using a MinElute PCR kit (Qiagen).

For *in vivo* screens, the “*In vivo Xenograft Model*” was used; T cells edited with the mini-pool library (**Supplementary Table 4**), editing was assessed by flow cytometry six days later, expansion continued for eight total days and then T cells were injected (n=4 mice, **Extended Data Fig. 5a**). Prior to injection, roughly 1e6 CAR T cells were frozen for subsequent gDNA isolation (NucleoSpin Blood kits). Ten days following T cell injection, mice were euthanized for tissue isolation. For bone marrow, long bones of each leg were isolated by dissection, crushed by mortar and pestle and 40 uM-strained. Extracted spleens were minced and gently 40 uM-strained. Red blood cells were lysed by incubating in ACK lysis buffer for two minutes followed by quenching with FACS buffer. Cells were resuspended in 400 uL FACS buffer and counted; 150 uL was used for flow cytometric analysis of cell populations and 250 uL was frozen for subsequent gDNA isolation (NucleoSpin Blood kits). For flow cytometric analysis, antibody mix was made in 100 uL FACS buffer and cells were stained for 30 minutes at 4°C with FcR-block (Miltenyi). Immediately before acquisition, cells were resuspended with 7-AAD at a 1:100 dilution and counting beads added.

For all screens, sequencing libraries were generated by amplicon sequencing using two rounds of PCRs with NEB Next Ultra II Q5 polymerase and primers purchased from IDT. The first PCR (98°C for 30”; 18 cycles of 98°C for 10”, 70°C for 30 s, 72°C for 45”; 72 °C for 2’) used a forward primer annealing to the “Internal Stuffer” with an Illumina Read 1 overhang and a reverse primer annealing to exon 1 of the *TRAC* locus with an Illumina Read 2 overhang (**Extended Data Fig. 2a)**. Following SPRI purification at a 1X volume ratio and input normalization, PCR (98°C for 30”; 12 cycles of 98°C for 10”, 60°C for 20”, 72°C for 25”; 72 °C for 2’) to append indexes and Illumina sequencing adaptors was performed using standard Illumina Nextera primers. Bead-purified, normalized and pooled libraries were sequenced on MiSeq or NextSeq instruments (Illumina).

Forward (**Bold**: Read 1 sequence): (5’**TCGTCGGCAGCGTCAGATGTGTATAAGAGACAG**CAACAATGTGCGGACGGCGT TG-3’)

Reverse (**Bold**: Read 2 sequence): (5’**GTCTCGTGGGCTCGGAGATGTGTATAAGAGACAG**TTGTTGCTCCAGGCCACAG CAC-3’)

### *In vivo* Xenograft Model

Eight-to-12-week old immunodeficient (NSG) mice purchased from Jackson Laboratory were dosed with 0.5e6 Ffluc-GFP Nalm6 via tail vein injection. Four days later, 0.1e6 CAR T cells were injected via the tail vein. One to two times per week, 200 uL (1.5 mg) of D-Luciferin Potassium Salt (Gold BioTechnology) was injected intraperitoneally and mice were imaged using an IVIS Spectrum Imaging System (Xenogen). Death was defined as hind limb paralysis, poor body score or loss of 15% or more of body weight. Experiments were carried out in accordance with a protocol approved by the University of California, San Francisco Institutional Animal Care and Use Committee. Analyses were performed in R using the ggsurvfit package.

### Genomics

#### Barcode Abundance Analysis

Each synthesized AP-1 domain contains a unique domain barcode (BC), with a minimum hamming distance of three between any two domain BCs. Unique DESynR TF BCs were constructed by concatenating the three constituent domain BCs with the “AGCG” cloning scar between each. In R, a reference of DESynR TF-BC associations was stored as a *PDict* (Biostrings package) object, raw fastq files were read in using *readFastq* (ShortRead package) and abundance was calculated using *vwhichPDict* (Biostrings package), allowing up to one mismatch. Log_2_FC was calculated by comparing DESynR BC-abundance in populations indicated in figure legends/main text using DESeq2^50^ default settings (design = ∼ Donor + Condition; where “Donor” indicates the human donor and “Condition” indicates the populations compared). Both raw counts and calculated Log_2_FCs are available for all screens (**Supplementary Table 2-4**). For the large DESynR AP-1 TF screen, analysis was filtered to constructs (n = 12,284) with at least 10 reads in input samples in all three human donors. For the mini-pool CD62L screen, “CD62L+” and “CD62L-” sorted populations were compared; CAR + GFP-control was omitted from analysis due to potential antibody fluorescence overlap with GFP signal. For mini-pool cytokine screens, cytokine positive sorted populations were compared to the “double-negative” population. For the mini-pool degranulation screen, “CD107a+” and “CD107a-” sorted populations were compared.

#### ATAC-Sequencing

Live CAR T cells (NGFR+) were sorted and nuclei were prepared from 50,000 cells per sample in duplicate by first washing in PBS + 0.04% BSA and then centrifugation in 50 ul lysis buffer (10 mM Tris-HCl pH 7.4, 10 mM NaCL, 3 mM MgCl_2_, 0.1% Tween-20, 0.1% IGEPAL, 0.01% Digitonin, 1% BSA in diH_2_O) for 10 minutes at 750g and 4°C. Nuclei were tagmented in 12.5 uL 2x TD Buffer, 10.5 uL diH_2_O and 2.5 uL Tn5 (Illumina) for 30 minutes at 37°C with agitation. Tagmented DNA was cleaned up using the DNA Clean & Concentrator-5 kit (Zymo Research), Nextera-indexed (Illumina) (72°C for 5’; 98°C for 30”; 12 cycles of 98°C for 10”, 63°C for 30”, 72°C for 1’) using NEB Next Ultra II Q5 polymerase and double-size selected using AMPure XP beads (Beckman Coulter). Libraries were normalized and pooled for sequencing on a NovaSeqX instrument with the following parameters: 51 bp Read 1, 12 bp i7 Index, 24 bp i5 Index, 51 bp Read 2.

ATAC-seq libraries were analyzed using PEPATAC with default settings^51^. Briefly, adapter sequences were trimmed with *--trimmer {skewer}* followed by removal of mitochondrial reads and reads associated with repetitive genomic regions. The remaining reads were aligned to the hg38 reference genome using Bowtie2^52^. Following alignment, SAMtools was used to extract uniquely mapped reads, and duplicate reads were removed using Picard generating final filtered BAM files for downstream processing. Peak calling for individual samples was performed with MACS2, and a consensus peak-set of non-overlapping 500-bp regions was generated^53^. The peak-sample count matrix was constructed using default settings of the *run_atac* function (ChrAccR; https://github.com/GreenleafLab/ChrAccR) and signal tracks for individual samples were merged using WiggleTools for group comparisons. The matrix was variance-stabilized using the *vst* function (DESeq2) and batch-effect corrected using *removeBatchEffect* in limma^54^. PCA plots were generated using the top 2,000 variable peaks and plotted using *plotPCA* (DESeq2). Differentially accessible peaks (abs(Log_2_FC)>0.5 & FDR<0.05) were identified with DESeq2. For heatmap visualization, differentially accessible peaks from all constructs relative to CAR + GFP control were aggregated, z-score standardized and subjected to k-means clustering. Additionally, construct-specific regulatory element enrichment was assessed using *matchMotifs* (motifmatchr; https://greenleaflab.github.io/motifmatchr/index.html) and with Homer motif position weight matrices (PWMs) from the gchromVAR package, which applies a weighting of chromatin features based on aggregated construct-specific peaks for comparison against random background peaks matched for GC content^55^.

#### scRNA-Sequencing

Live CAR T cells (NGFR+) were sorted from each unique sample and stained with MULTI-seq barcodes (Sigma Aldrich). Stained cells were pooled in equal proportions and 25,000 cells per lane were loaded onto a Chromium Controller (10X). 5’ gene expression libraries were prepared per manufacturer instructions and MULTI-seq libraries were prepared as previously described^56^. Gene expression and MULTI-seq libraries were quantified using an Agilent Bioanalyzer and pooled 90:10 for sequencing on a NovaSeqX instrument with the following parameters: 28 bp Read 1, 10 bp i7 Index, 10 bp i5 Index, 91 bp Read 2.

Fastqs for each sample were processed with *cellranger count* version 6.0.0 with the GRCh38 v3.0.0 transcriptome and then aggregated together using *cellranger aggr*^57^. MULTI-seq tags–which indicate the construct each cell was edited with and the human donor each cell came from–were assigned to cells using *deMULTIplex2*^56,58^. Following tag assignment, predicted multiplets and low confidence assignments were filtered from downstream analysis. The resulting cell by gene matrix and assignment metadata was further processed using Seurat v5.1.0 and *SeuratObject* v5.0.2. A 20-component PCA was computed on normalized and scaled data using the top 2,000 variable features^59–63^. To regress donor-specific differences, harmony v1.2.0 was called using the *IntegrateLayers* function within Seurat and the original PCA^64^. A small number of Nalm6 cells expressing B cell marker genes were then identified and removed from downstream analysis and the initial normalization, PCA, and integration were redone. Unsupervised clusters were called using the integrated dimension reduction and used for cell type assignment. Assignment was performed based on expression of top differentially expressed genes (DEGs) identified with Wilcoxon rank sum testing using the ‘*Presto*’ implementation^65^. Positive markers for each donor and construct pair were identified against control CAR + GFP control cells from the same donor using 500 randomly selected cells from each category to compare numbers of DEGs across samples of variable depth. Only genes with abs(Log_2_FC)>0.5 and >5% expression in at least one of the two categories were considered in DEG testing.

#### Pseudobulk of scRNA-Sequencing Data

Pseudobulk profiles of scRNA-seq data by construct and donor were generated by extracting raw counts from the Seurat object and calling the *aggregate.Matrix()* function (Seurat). The count matrix was variance-stabilized with the *vst* function (DESeq2) and batch corrected with *removeBatchEffect* (limma). PCA plots were generated with the *plotPCA* function (DESeq2) using the top 1,000 most variable genes. Differentially expressed genes (abs(Log_2_FC)>0.5 & FDR<0.05) were identified with DESeq2. For heatmap visualization, differentially expressed genes from all constructs relative to CAR + GFP control were aggregated, z-score standardized and subjected to k-means clustering.

#### Permutation Testing

Permutation testing of the correlation between DEGs and DARs in corresponding Modules (Fig. 4h**,g**) was performed using ChIPpeakAnno and regioneR (R packages)^66,67^. Briefly, nearest genes were assigned to each DAR with *annoPeaks* function (ChIPpeakAnno). Then, the *permTest* function (regioneR) was used to test the overlap between annotated genes and those in the linked DEG Module. The number of overlapping genes between each linked DAR and DEG Module was tested against a background of overlaps between DAR-associated genes and 1,000 permutations of a randomly sampled gene-set (all DEGs) of the same size as the DEG Module in question.

#### Long-Read Sequencing

From the HDRT library, an amplicon spanning upstream *TRAC* homology, tNGFR, CAR, TF and downstream *TRAC* homology was PCR-amplified (98°C for 30”; 15 cycles of 98°C for 10”, 60°C for 15”, 72°C for 2’; 72 °C for 2.5’) using NEB Next Ultra II Q5 polymerase and cleaned up using 1X AMPure PB beads (Pacific Biosciences). The sequencing library was prepared using the SMRTBell Library Prep Kit 3.0 (Pacific Biosciences) following manufacturer’s instructions. The final library was quantified using a dsDNA HS kit on a Qubit instrument (Thermo Fisher) and using Genomic DNA ScreenTapes on a Tapestation 4200 instrument (Agilent) and sequenced on a Revio instrument (Pacific Biosciences).

Forward:

5’-GGAACCGGTGCTGGAAGTGGT-3’

Reverse:

5’-CACTGAGCCTCCACCTAGCCT-3’

Domain barcodes and sequences were extracted from each PacBio HIFI read using the following procedure. Each of the 11-bp domain BCs were concatenated with the AGCG cloning scar sequence to decrease the likelihood of identifying an off-target BC-match at a different read locus. The resulting sequences and their reverse complements were identified in each read by a brute force search for perfect matches and saved along with the associated read. Three “references” were constructed to align reads against, containing N domain, bZIP domain, and C domain sequences. The bowtie2 aligner was used to identify sequence matches with *--very-sensitive-local* alignment parameters. The top scoring alignment for each domain and the associated read was extracted for downstream analysis. Manual scanning of reads before filtering revealed that most cases of missing domains based on bowtie2 alignments were the result of unsuccessful alignment of reads to short domains. A cutoff of 50 bp was chosen and 6 out of 72 domains were removed from downstream analysis. Following domain and BC assignment, each read was iterated over and assigned one classification, including “correct reads”, “domain mismatches”, “missing domains”, and “missing BCs”.

## Data Availability

All scRNA-seq and bulk ATAC-seq data have been deposited to NIH GEO under accession numbers, GSE280508 and GSE280509. Long-read sequencing data is available upon request.

## Code Availability

No customs algorithms were developed, and bioinformatics tools used are described in the Methods. Code is available upon request.

## Acknowledgements

We thank the members of the Satpathy Lab for stimulating discussions. A.T.S. was supported by a Career Award for Medical Scientists from the Burroughs Wellcome Fund, a Lloyd J. Old STAR Award from the Cancer Research Institute, and the Parker Institute for Cancer Immunotherapy. This work was supported by the Department of Pathology at Stanford University. Sequencing was performed at the UCSF CAT, supported by UCSF PBBR, RRP IMIA, and NIH 1S10OD028511-01 grants. The authors thank the DNA Technologies and Expression Analysis Core at the UC Davis Genome Center, supported by NIH Shared Instrumentation Grant 1S10OD010786-01.

## Author Contributions

O.T., T.L.R., and A.T.S conceptualized the study. O.T., T.L.R., and A.T.S. wrote and edited the manuscript with input from all authors. O.T., Y.Y., G.C.R., C.K., J.L., A.K.M., C.J.R., N.E.T, and P.K.Y. performed experiments. A.H. and A.Y.C. set up computational pipelines and analyzed the data. J.E., T.L.R., and A.T.S. guided experiments and data analysis.

---

**Extended Data Figure 1:**
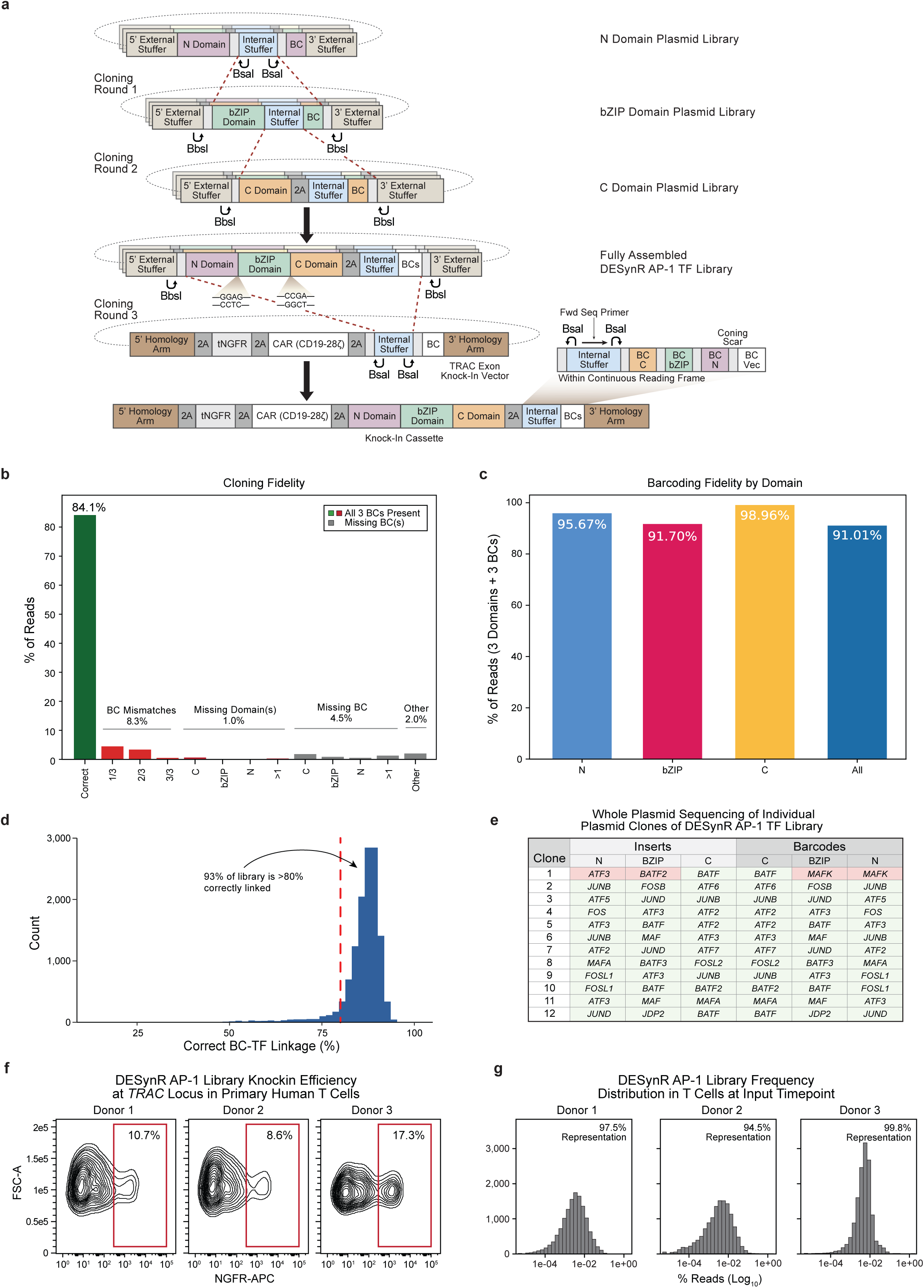
Construction and quality control of DESynR libraries. **a)** Libraries of full-length DESynR genes were iteratively constructed from individual domains and subcloned into a CD19-28ζ-CAR- and tNGFR-reporter-containing vector for primary human T cell screening. Each domain was synthesized with a constant Type IIS restriction enzyme-excisable region (“Internal Stuffer”) followed by a unique 11-base pair (bp) barcode (BC) and flanked by additional constant external adapter sequences on the 5’ and 3’ ends containing distinct Type IIS restriction enzyme cut sites (“5’ External Stuffer” & “3’ External Stuffer”). Three domain pools were formed from individually synthesized members: “N”, “bZIP” and “C”. To append the “bZIP” domain to the “N” domain, the “bZIP” domain was cleaved externally (BbsI) and the “N” domain was cleaved internally (BsaI) and then ligated. The “C” domain was then appended to the “N+bZIP” domain-pair analogously. Domains were not separated by linker sequences so minor sequence modifications affecting one to two amino acids were made to standardize the four-bp overhangs (see **Supplementary Data Table 1**). The 3’ cloning scar separating each domain BC was constant (“AGCG”). **b-d,** Long-read sequencing of the final plasmid library, where each read spanned a single fully assembled DESynR AP-1 TF and its BC region to assess barcoding fidelity and cloning efficiency. **b)** A bar plot considering all sequencing reads. “All 3 BCs Present” indicates reads from constructs that contained three valid BCs and thus contributed to BC-abundance calculations from next-generation sequencing (NGS) data. The low-frequency reads captured by “Missing BC” and “Other” had less than three BCs and were not considered in abundance analysis **c)** A bar plot of overall (all constructs aggregated) domain-BC linkage fidelity from reads containing a full three-domain DESynR AP-1 TF and three BCs (“Correct” and “BC Mismatches” in **b**). Over 90% of assembled DESynR AP-1 TFs had a correct linkage of BCs across all three domains. **d)** A histogram of correct TF-BC linkage frequency; for each DESynR TF defined by a unique combination of three domains, the frequency of reads where all domains and BCs matched was considered correct TF-BC linkage. Over 90% of DESynR TFs had high (>80%) fidelity. **e)** A table summarizing an orthogonal whole plasmid sequencing analysis of TF-BC linkage with resolution of each domain from individual clones (n=12), showing similarly high fidelity. **f)** Flow cytometry of editing efficiency with the DESynR AP-1 TF library in primary human T cells (n=3 healthy donors). **g)** Histograms of library distribution in edited primary human T cells read out by NGS of BCs (n=3 healthy donors).

**Extended Data Figure 2:**
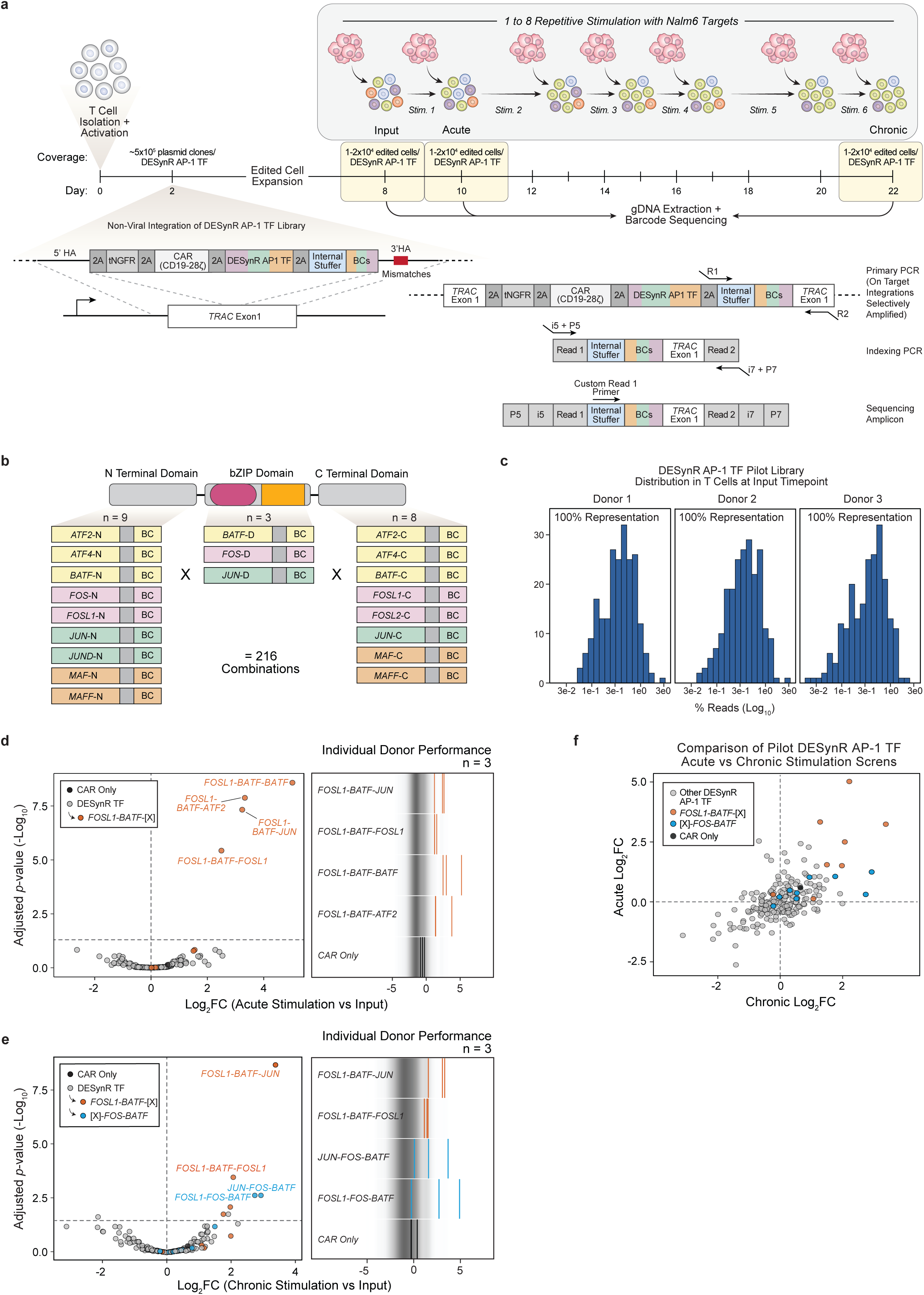
Development of an *in vitro* screening platform for DESynR AP-1 TF-expressing CAR T cells. **a)** A timeline schematic of the pooled screening platform. Primary human T cells from healthy donors were activated with CD3/28 agonism. Two days later, Cas9 ribonucleoprotein targeting exon 1 of *TRAC* was electroporated into T cells along with the CAR + DESynR AP-1 TF plasmid library as a homology-directed repair template; this resulted in in-frame integration within the first exon of *TRAC*, enabling transcription of the CAR, tNGFR reporter and DESynR AP-1 TF from the endogenous *TRAC* promoter^32,33^. Following six days of expansion, the T cells were subjected to an *in vitro* repetitive stimulation assay. Every two-to-three days for six total stimulations over two weeks, counting beads and flow cytometry were used to determine how many CAR T cells and Nalm6 targets were in culture and the CAR T cell to Nalm6 ratio was standardized to one to eight. Genomic DNA (gDNA) was extracted from cells prior to the assay (“Input”), following one stimulation (“Acute”) and following six stimulations (“Chronic”). The BC region was amplified from gDNA using a forward primer that annealed directly upstream of the BC region within the “Internal Stuffer” and a reverse primer that annealed downstream in *TRAC* Exon 1. Several mismatches were included in the 3’ *TRAC* homology arm of the repair templates at the binding site of the reverse primer, such that only on-target integrations were selectively amplified^22^. A secondary indexing PCR produced an Illumina-compatible, sequencing-ready amplicon. Coverage estimates of the library from plasmid to edited cells and through the assay are indicated. **b)** Schematic of the natural AP-1 TF domain pools used to generate a small pilot library for validation of the screening pipeline. **c)** Histograms of the DESynR AP-1 TF pilot library distribution in edited T cells at the “Input” timepoint read out by NGS of barcodes (n=3 human donors). **d-f,** Log_2_FC calculations of BC abundance between “Acute” and “Input” and “Chronic” and “Input” timepoints were performed with DESeq2; significance was calculated by the Wald test and corrected for multiple comparisons using the Benjamini-Hochberg method (n=3 human donors). **d)** Left, a volcano plot indicating the enrichment or depletion of each construct following acute stimulation. Hashed horizontal line indicates *p*=0.05. Right, Log_2_FC in individual human donors for selected constructs following acute stimulation. Each vertical bar represents one human donor. **e)** Left, a volcano plot indicating the enrichment or depletion of each construct following chronic stimulation. Hashed horizontal line indicates *p*=0.05. Right, log_2_FC in individual human donors for selected constructs following chronic stimulation. Each vertical bar represents one human donor. **f)** Scatter plot comparing Log_2_FC in acute and chronic stimulation screens. A *FOSL1-BATF* “N+bZIP” domain-pair signature was strongly enriched in both acute and chronic stimulation screen top-performers, while a *FOS-BATF* “bZIP + C” signature was enriched specifically in the chronic stimulation screen.

**Extended Data Figure 3:**
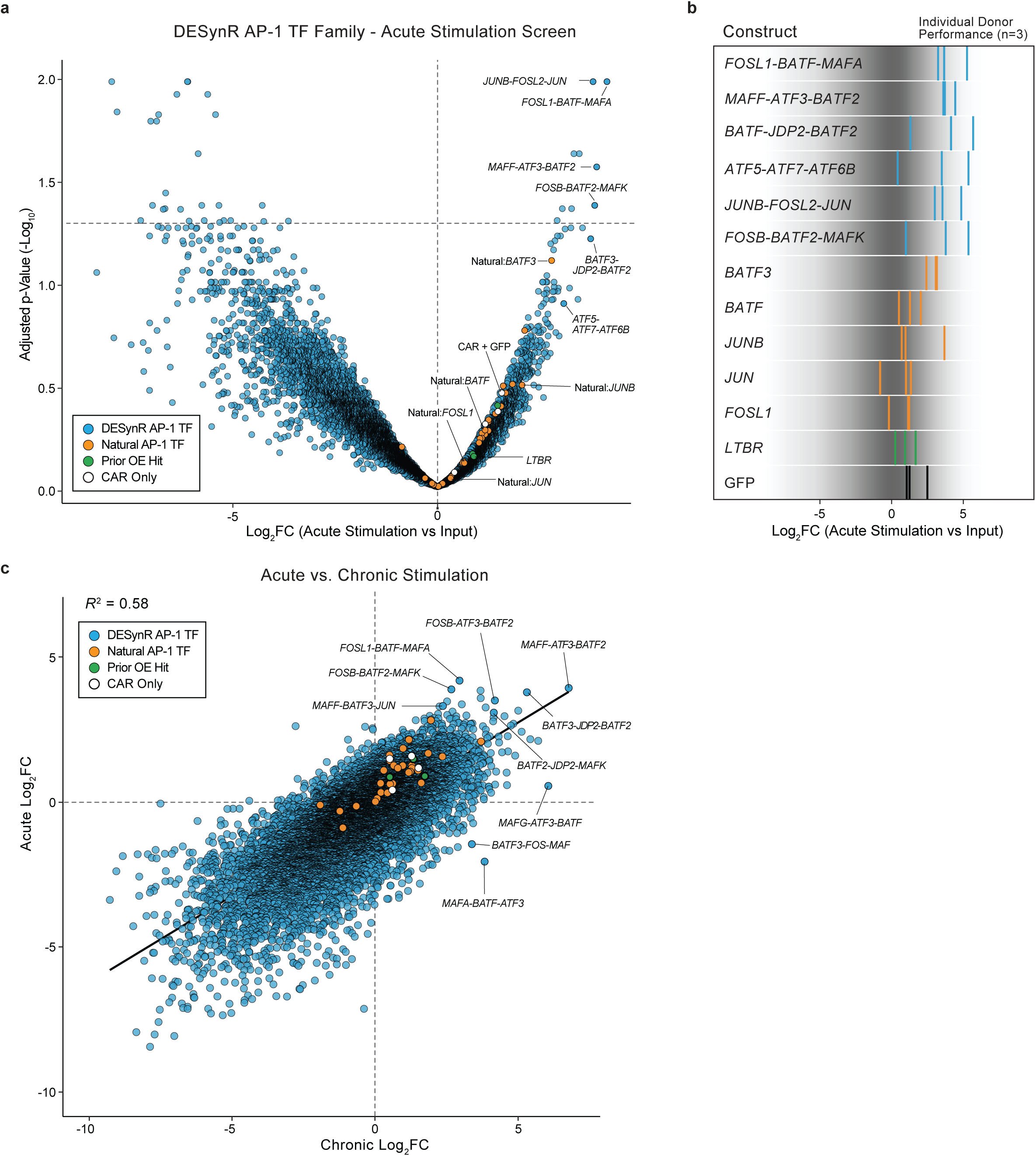
DESynR AP-1 TFs outperform natural AP-1 TFs in an acute stimulation CAR T cell screen. **a-c**, Log_2_FC calculations were performed with DESeq2; significance was calculated by the Wald test and corrected for multiple comparisons using the Benjamini-Hochberg method (n=3 human donors). **a)** A volcano plot indicating the enrichment or depletion of each DESynR AP-1 TF (blue), natural AP-1 TF (orange), previous overexpression candidates (e.g. *LTBR*; green) and CAR-only controls (white) following acute stimulation (**Extended Data** Fig. 2). Hashed horizontal line indicates *p*=0.05. **b)** Log_2_FC in individual human donors for selected constructs following acute stimulation. Each vertical bar represents one human donor (n=3). **c)** Scatter plot comparing Log_2_FC in acute and chronic stimulation screens. R^2^ calculated from a y ∼ x linear regression.

**Extended Data Figure 4:**
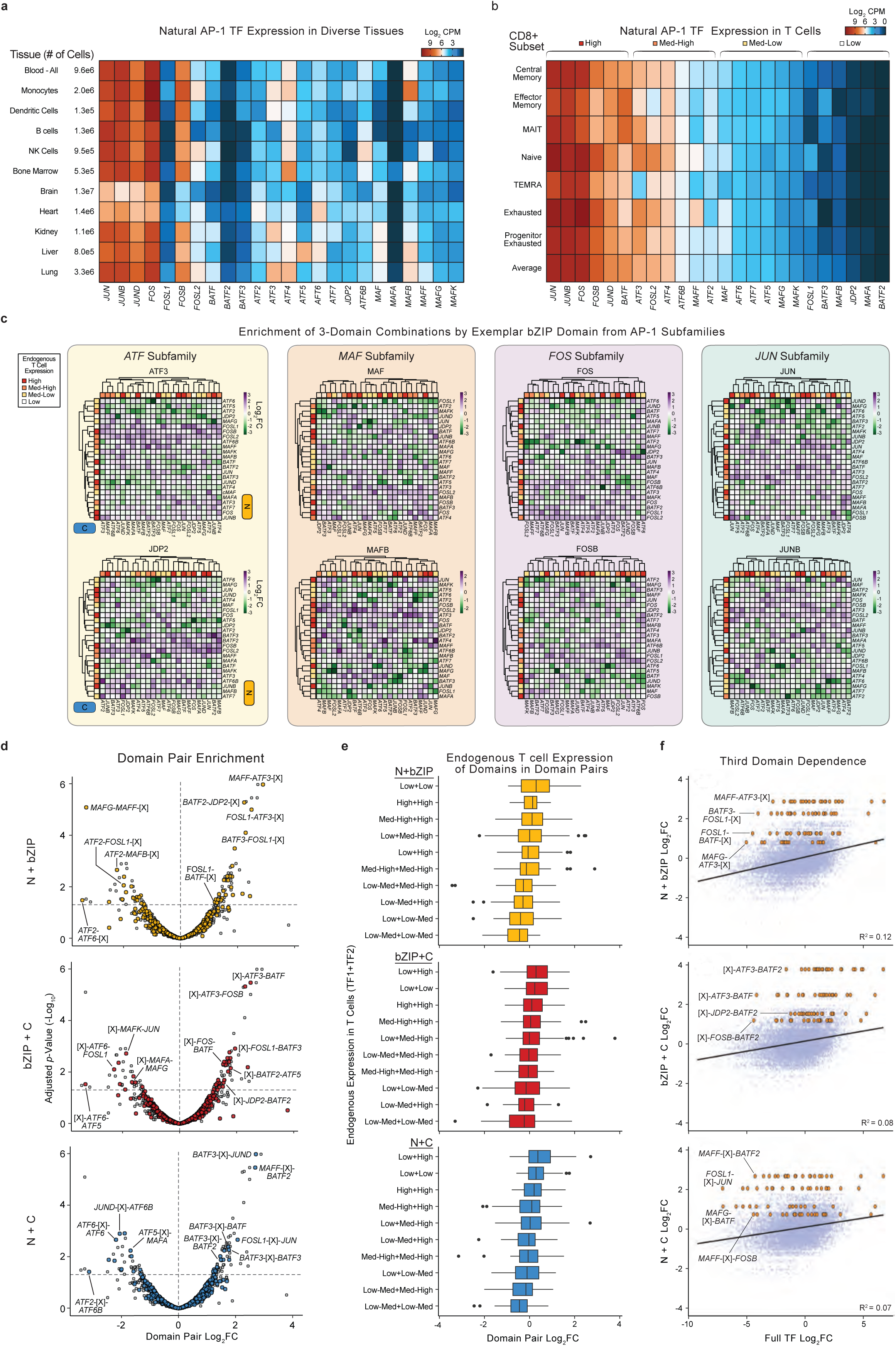
Domain-use syntax of top DESynR AP-1 TFs. **a)** A heatmap of average expression of 24 natural AP-1 TFs across various tissues. Data from the Chan-Zuckerberg Initiative Cell x Gene Portal^47^. **b)** A heatmap of average expression of 24 natural AP-1 TFs across CD8 T cell subsets^48^. The quartile annotation of expression of natural AP-1 TFs in endogenous T cells (**c,e** and Fig. 2e**,f**) is highlighted. **c)** Heatmaps of Log_2_FC for “N+C” domain-pairs for two example “bZIP” domains from each of the four AP-1 subfamilies. Each cell of the heatmap represents a unique “N+C” domain-pair. Outside heatbars report expression of the domain’s corresponding full natural AP-1 TF in endogenous T cells (colored by quartile). **d-f,** Each three-domain DESynR AP-1 TF was re-classified as three separate domain-pair architectures (“N+bZIP”, “bZIP+C” and ”N+C”). Log_2_FC calculations were performed with DESeq2; significance was calculated by the Wald test and corrected for multiple comparisons using the Benjamini-Hochberg method (n=3 human donors). **d)** Volcano plots indicating the enrichment or depletion of each domain-pair following chronic stimulation. Hashed horizontal line indicates *p*=0.05. **e)** Boxplots of domain-pair Log_2_FC by expression (split into quartiles: Low, Low-Med, Med-High, High) of constituent domains in endogenous T cells. Annotations correspond to domains (e.g. “Low-High” in “bZIP-C” indicates a lowly expressed “bZIP” domain and a highly expressed “C” domain). Each architecture is shown separately and organized in descending order of Log_2_FC. Across domain-pairs, exclusive utilization of domains from lowly expressed natural AP-1 TFs, as well as one lowly and one highly expressed domain, showed the highest enrichment. **f)** Scatter plots of Log_2_FC of domain-pairs versus Log_2_FC of three-domain TFs (from Fig. 2b) with each architecture shown separately. A given domain-pair appears up to 24 times with a unique third domain as a three-domain DESynR AP-1 TF. Select domain-pairs are highlighted, demonstrating that even within enriched domain-pairs, the identity of the third domain can have a large influence on the overall functional performance of the three-domain DESynR AP-1 TF. R^2^ calculated from y ∼ x linear regressions.

**Extended Data Figure 5:**
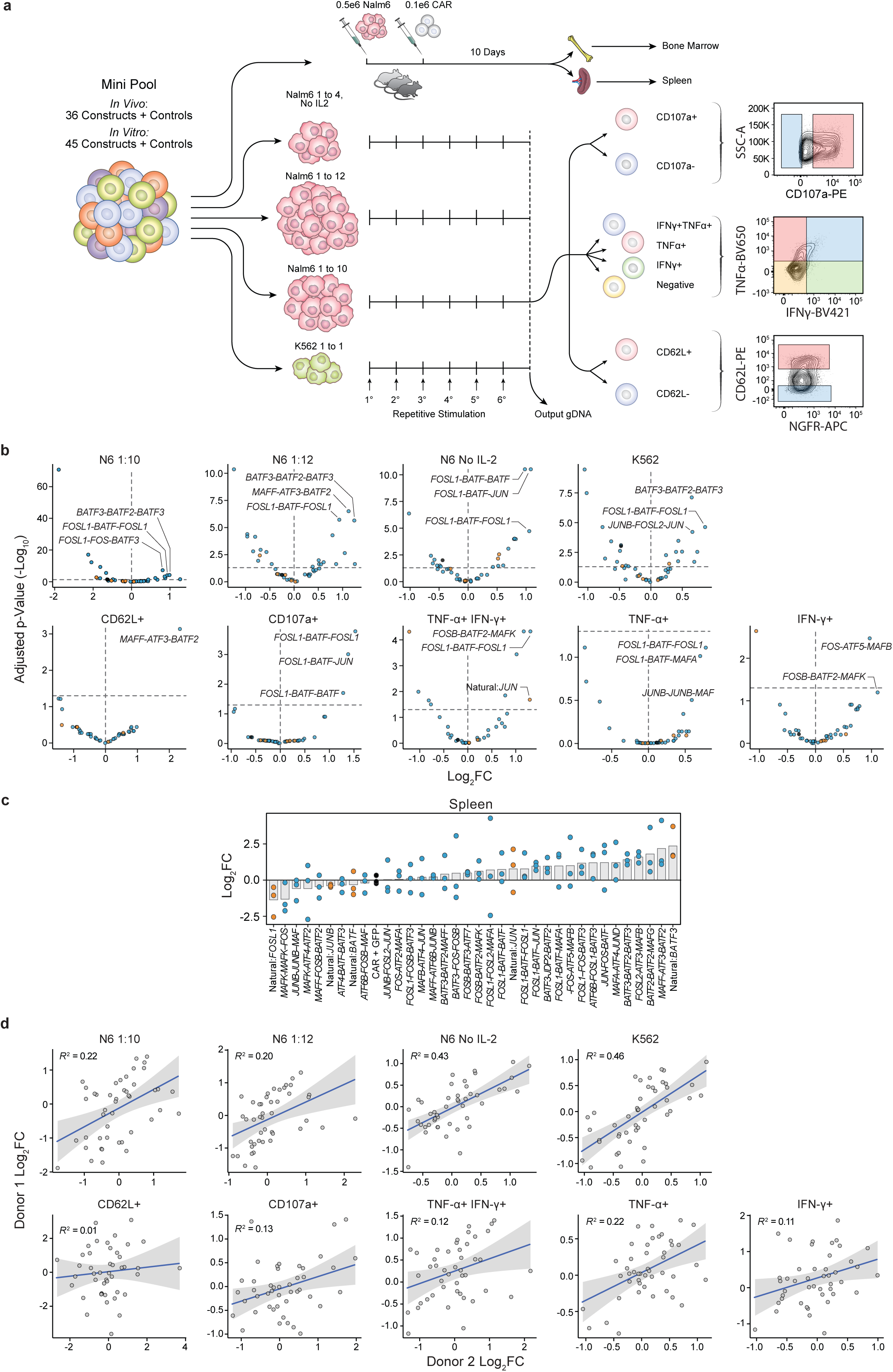
*In vitro* and *in vivo* functional screening of a DESynR AP-1 TF mini-pool. **a)** Schematic of mini-pool functional screening. For repetitive stimulation assays, a single input sample (edited CAR T cells pre-assays) served as the comparator for all output (post-assays) samples (n=3 human donors). Chronically stimulated (six total stimulations; every two-three days for two total weeks) cells from the “Nalm6 1 to 10 (Med)” assay were used for cytokine (IFNγ/TNFα) production, degranulation (CD107a) and CD62L-expression sorts (n=2 human donors). The colored boxes indicate FACS gates; for all FACS-based assays, the sorted negative population served as the comparator. For *in vivo* screens, CAR T cells from each organ (spleen/bone marrow) harvested ten days post-CAR T cell injection were compared to a single input (edited CAR T cells pre-injection) sample (n=1 human donor, 3-4 mice). **b)** Volcano plots of each individual screen. Log_2_FC calculations were performed with DESeq2; significance was calculated using the Wald test and corrected for multiple comparisons using the Benjamini-Hochberg method. Horizontal dashed line indicates *p*=0.05. Select TFs are annotated. **c)** Bar plot of Log_2_FC in abundance in the spleen from the *in vivo* Nalm6 xenograft screen scaled to CAR + GFP-control (black). Each overlaid point represents one mouse (n=3). **d)** Scatter plot of the correlation in Log_2_FC in two representative human donors across all *in vitro* screens (n=2-3 human donors). R^2^ calculated from y ∼ x linear regressions.

**Extended Data Figure 6:**
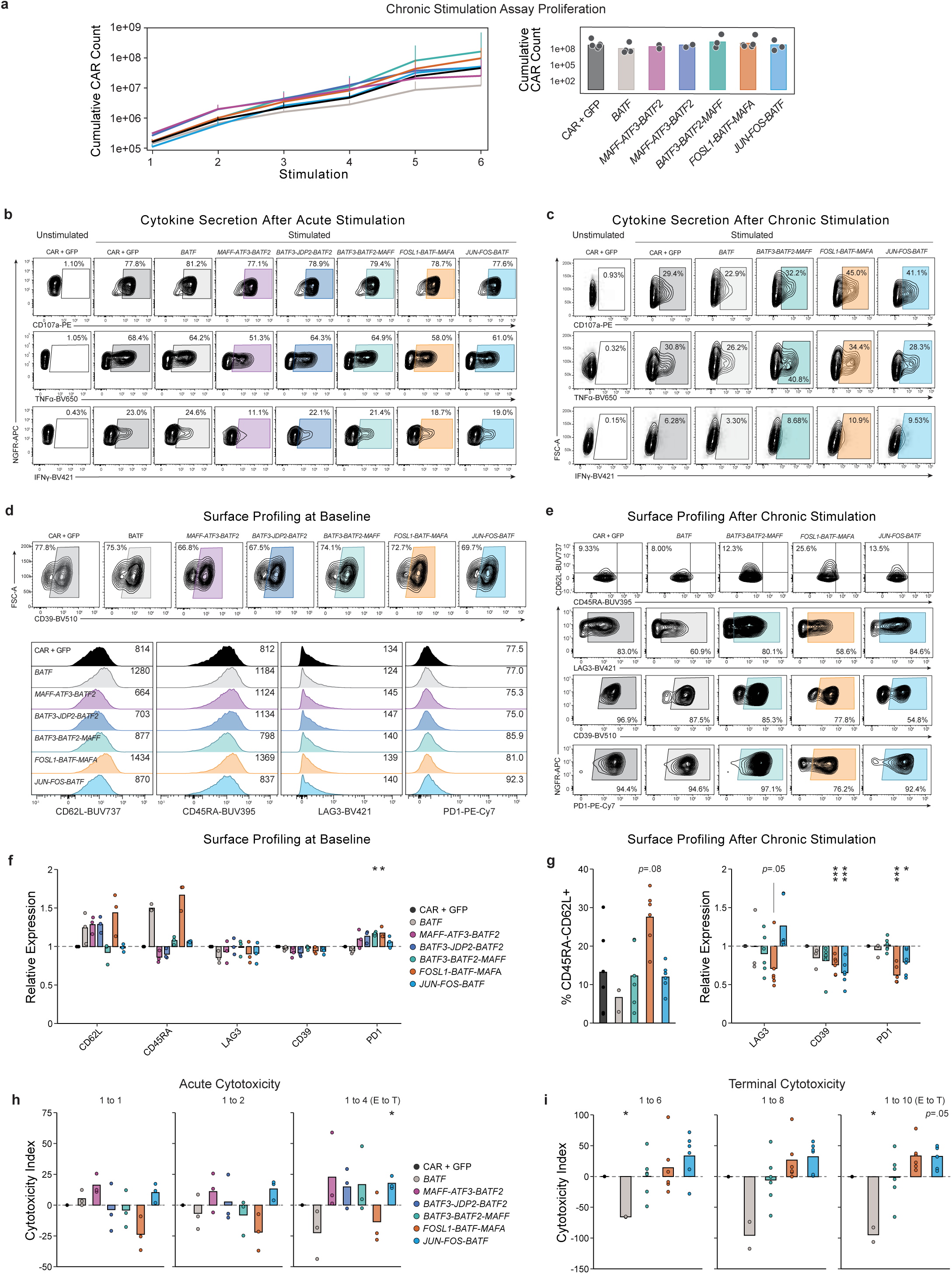
Individual *in vitro* validation of top DESynR AP-1 TFs. **a)** Proliferation in an arrayed two-week chronic stimulation assay with Nalm6 target cells (n=2-6 human donors). Left, a line graph of mean count +/-s.e.m. at each stimulation. Right, a bar plot of cumulative count following six stimulations. **b)** Exemplar flow cytometry from one representative human donor of degranulation and cytokine production at baseline following a single acute stimulation with Nalm6 target cells (CAR T cells not previously stimulated; n=2-5 donors). **c)** Exemplar flow cytometry from one representative human donor; cells were stimulated a final time to measure degranulation and cytokine production following a two-week chronic stimulation assay (n=2-8 donors). **d)** Exemplar flow cytometry from one representative human donor of key surface proteins at baseline (CAR T cells not previously stimulated; n=2-6 donors). **e)** Exemplar flow cytometry from one representative human donor of key surface proteins following a two-week chronic stimulation assay (n=2-6 donors). **f)** Bar plots of expression of key surface proteins at baseline (CAR T cells not previously stimulated). Overlaid points represent average of technical replicates for one human donor (n=2-3). Significance calculated by a paired two-tailed Student’s *t*-test. **g)** Bar plots of expression of key surface proteins following a two-week chronic stimulation assay. Overlaid points represent average of technical replicates for one human donor (n=2-6). Left, raw expression (see gating top row in **e**). Right, expression relative to CAR + GFP-control. Significance calculated by a paired two-tailed Student’s *t*-test. **h)** Bar plots of *in vitro* acute cytotoxicity of Nalm6 target cells (CAR T cells not previously stimulated). Cytotoxicity index defined as cytotoxicity relative to CAR + GFP-control determined from a flow cytometric count of remaining Nalm6 target cells. Overlaid points represent average of technical duplicates or triplicates in n=3 human donors. Significance calculated by a paired two-tailed Student’s *t*-test. **i)** Bar plots of cytotoxicity in a terminal challenge following two weeks of chronic stimulation. Overlaid points represent average of technical duplicates or triplicates (n=2-6 human donors). Significance calculated by a paired two-tailed Student’s *t*-test. \**p*<0.05, \*\**p*<0.01, \*\*\**p*<0.0001

**Extended Data Figure 7:**
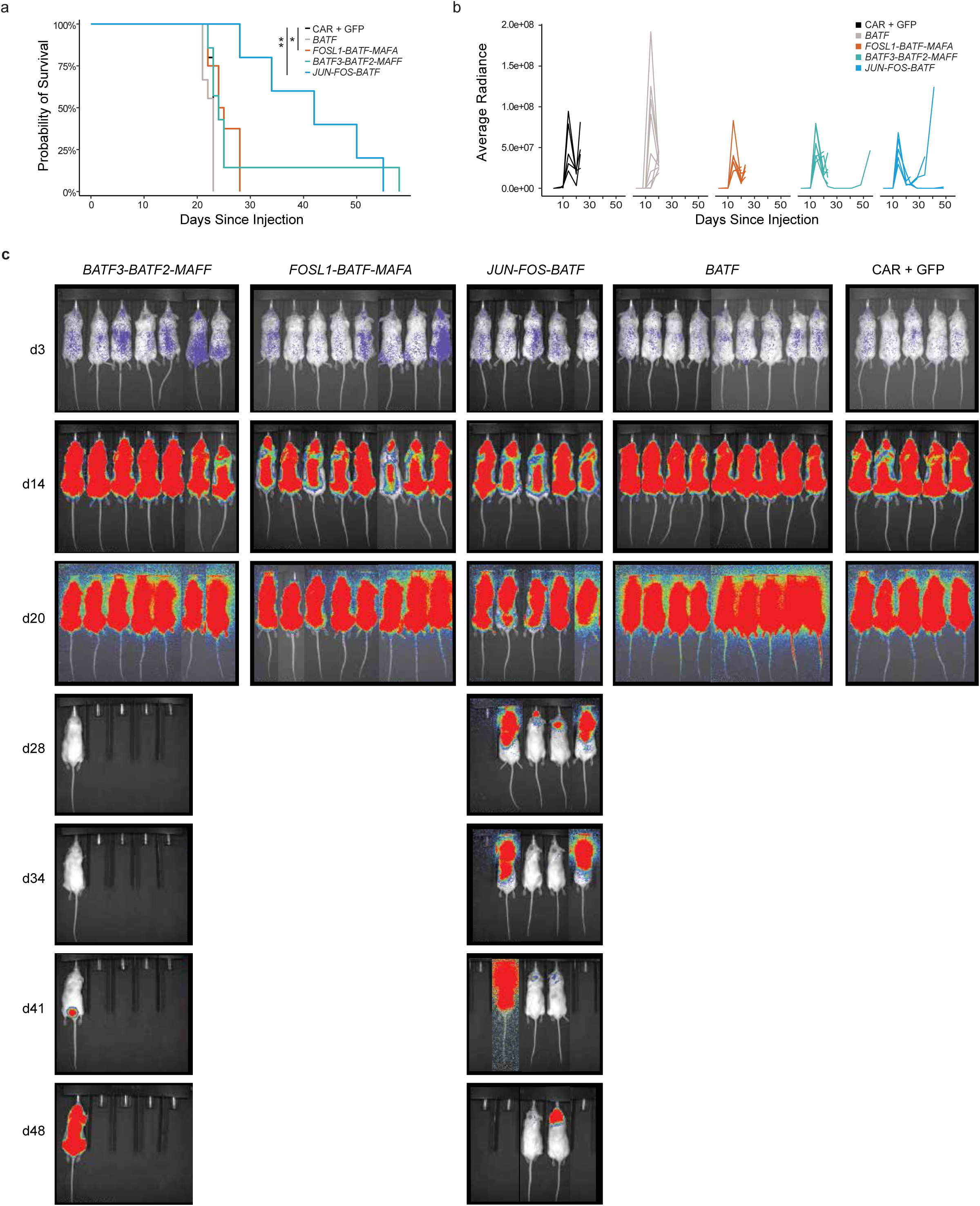
Functional *in vivo* validation of top DESynR AP-1 TFs. **a-c**, 0.5e6 Nalm6 human leukemia cells were injected intravenously (IV) into immunodeficient mice, followed four days later by IV-injection of 0.1e6 human CAR T cells edited with indicated DESynR AP-1 TFs, natural *BATF* or CAR + GFP-control. Mice were followed to endpoint (n=5-9). **a)** A Kaplan-Meier survival curve. Significance of each construct versus CAR+GFP-control was calculated by a Mantel-Cox test. \**p*<0.05, \*\**p*<0.01. **b)** A line graph of tumor burden quantified by bioluminescence imaging with each construct shown separately. **c)** Raw bioluminescence images from selected timepoints.

**Extended Data Figure 8:**
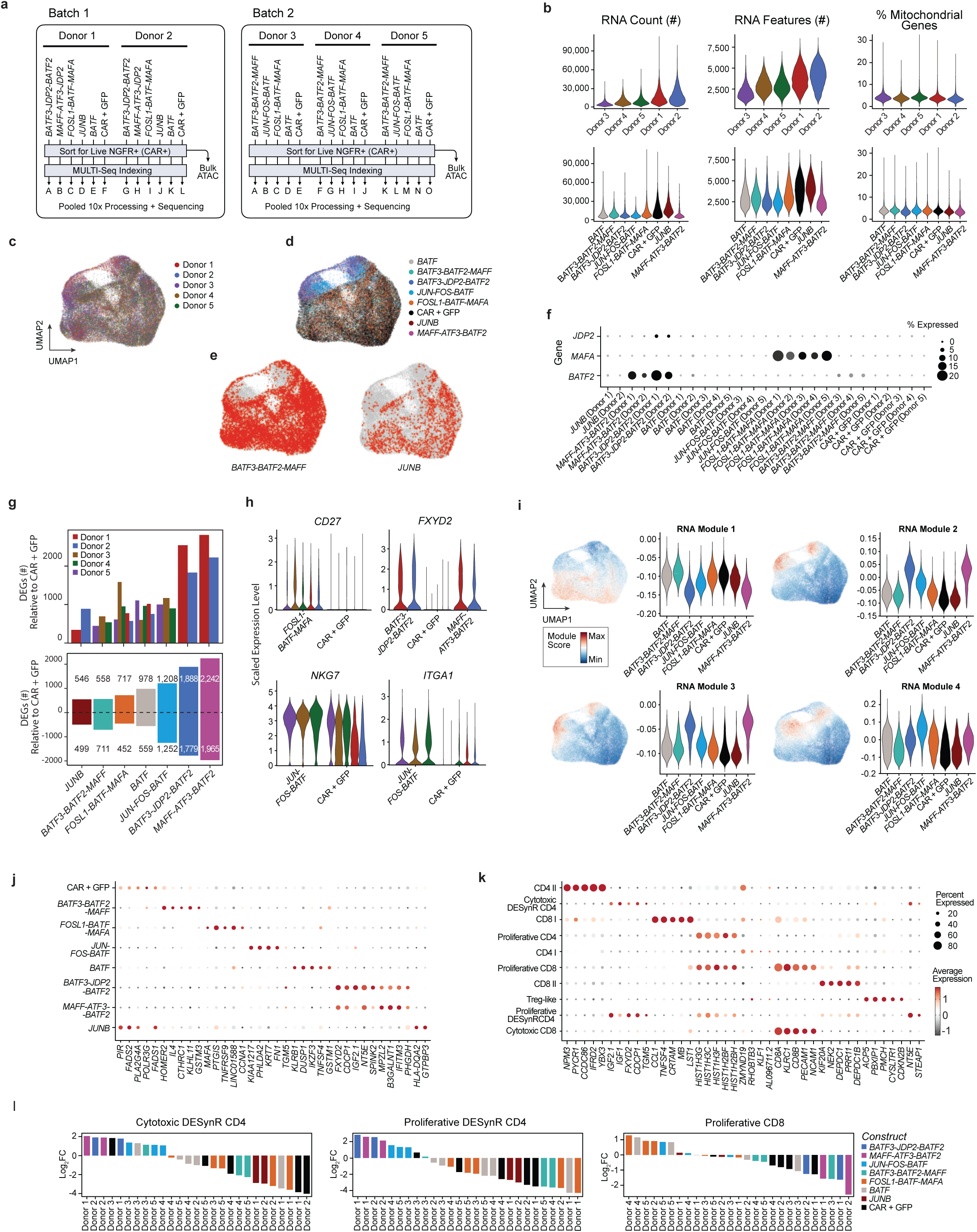
Characterization of DESynR and natural AP-1 TFs by single-cell RNA-sequencing. **a)** A schematic overview of sample processing for scRNA-seq and bulk ATAC-seq experiments (n=7 constructs, 5 human donors, 2 batches). **b)** Violin plots of scRNA-seq quality control metrics by human donor (top) and construct (bottom). **c)** Uniform manifold approximation and projection (UMAP) of scRNA-seq data colored by human donor (n=5) shows intermixing of cells from the five tested human donors following integration. **d)** UMAP of scRNA-seq data colored by construct. **e)** UMAPs of scRNA-seq data with red dots indicating cells of selected constructs and all other cells as gray dots (additional tested constructs shown in Fig. 4c). **f)** A dot plot indicating percent expression (size) of mRNA transcripts from TF constructs as algorithmically assigned from scRNA-seq data. Individual domains included within the DESynR AP-1 TFs were likely detected within the expressed TCRα mRNA transcript and misclassified as an mRNA transcript from the natural AP-1 TF from which the domain was derived. **g)** Bar plots of differentially expressed genes (DEGs) from scRNA-seq data. Top, all DEGs split by human donor and construct. Bottom, DEGs averaged from integrated donors for each construct. Shown in ascending order of total DEGs with count of increased and decreased genes indicated. **h)** Violin plots of exemplar marker genes by construct and human donor in scRNA-seq data. **i)** Pseudobulk RNA-seq data was hierarchically clustered into four gene modules (Fig. 4g). Expression of these modules on the UMAP of scRNA-seq data (left) and violin plots of scaled expression by construct in scRNA-seq data (right). Module 4 expression is uniquely confined to areas of the UMAP occupied by *JUN-FOS-BATF*-expressing cells, although original clustering did not define a single cluster in this region. (Fig. 4a**-d**) **j)** A dot plot of the percentage of cells expressing (size) and scaled average expression (color) for select marker genes by construct. **k)** A dot plot of the percentage of cells expressing (size) and scaled average expression (color) for select marker genes in manually annotated clusters in UMAP of scRNA-seq data. **l)** Bar plots of Log_2_FC in abundance of constructs in selected clusters of the UMAP of scRNA-seq data split by donor.

**Extended Data Figure 9:**
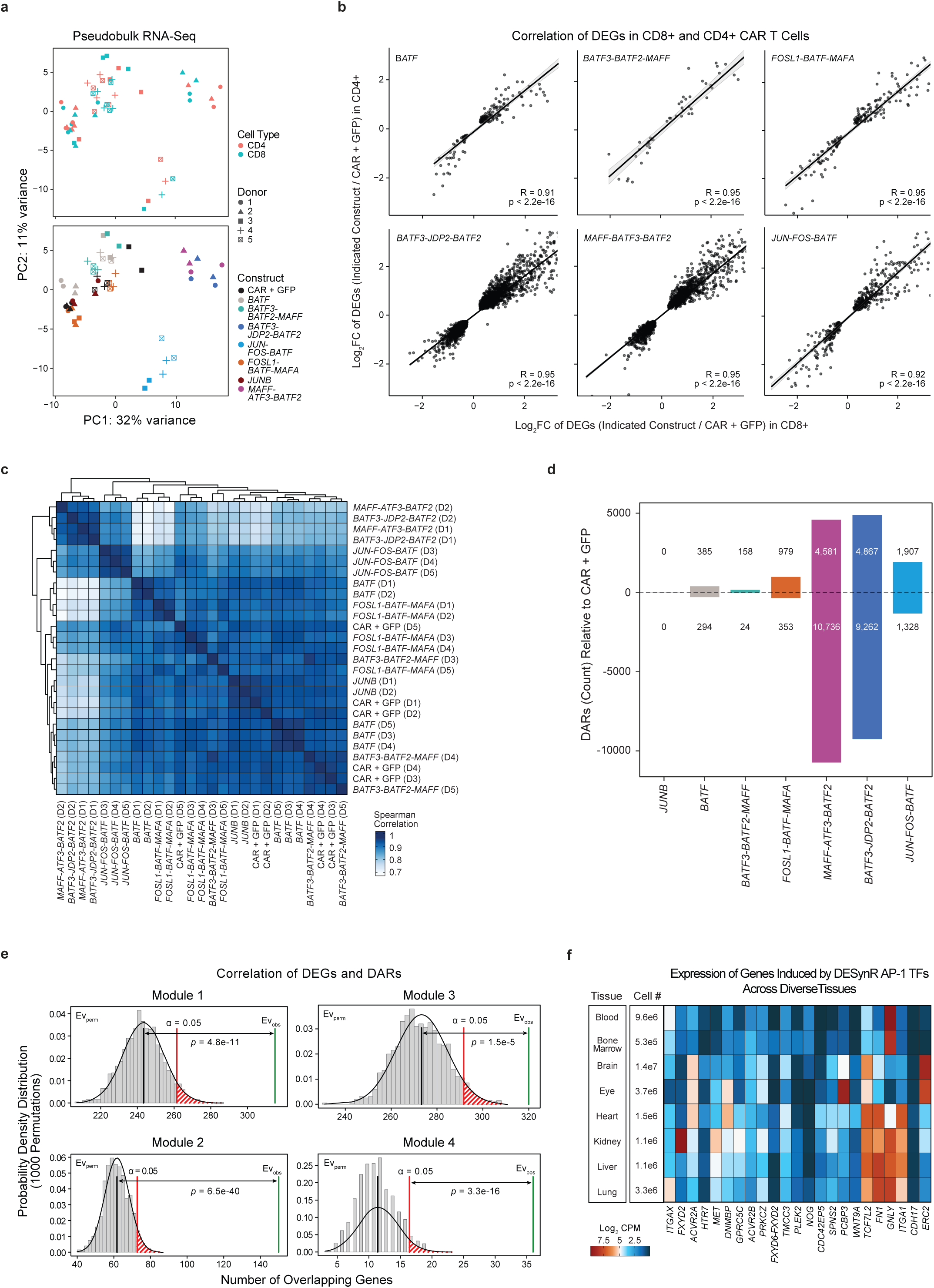
Characterization of DESynR and natural AP-1 TFs by bulk transcriptional and epigenetic profiling. **a)** Principal component analysis (PCA) of pseudobulk RNA-seq data. Top, colored by T cell subset (CD4+ vs. CD8+) and a unique shape for each human donor (n=5). Bottom, colored by construct and a unique shape for each human donor (n=5). **b)** Scatter plots of Log_2_FC of DEGs in CD4+ versus CD8+ cells for six of eight constructs from pseudobulk RNA-seq data. R^2^ calculated from y ∼ x linear regressions and *p*-values from Pearson’s correlation. **c)** Heatmap reporting Spearman correlation between individual samples (construct and human donor delineated) of bulk ATAC-seq data. All peaks considered. **d)** Bar plot of differentially accessible chromatin regions (DARs) relative to CAR + GFP-control for each construct from bulk ATAC-seq data. Direction (increased or decreased) is indicated, and magnitude (number of regions) is annotated. **e)** Histograms of results from permutation tests linking chromatin accessibility and gene expression changes within each module. For each module, gene sets were permuted 1000 times to test whether chromatin regions were significantly enriched near genes in that specific module compared to a random background gene set. Each module is shown separately. **f)** A heatmap of average expression of selected DEGs (from pseudobulk RNA-seq data) across different tissues. Tissue expression data from the Chan-Zuckerberg Initiative Cell x Gene Portal^47^.

**Extended Data Figure 10:**
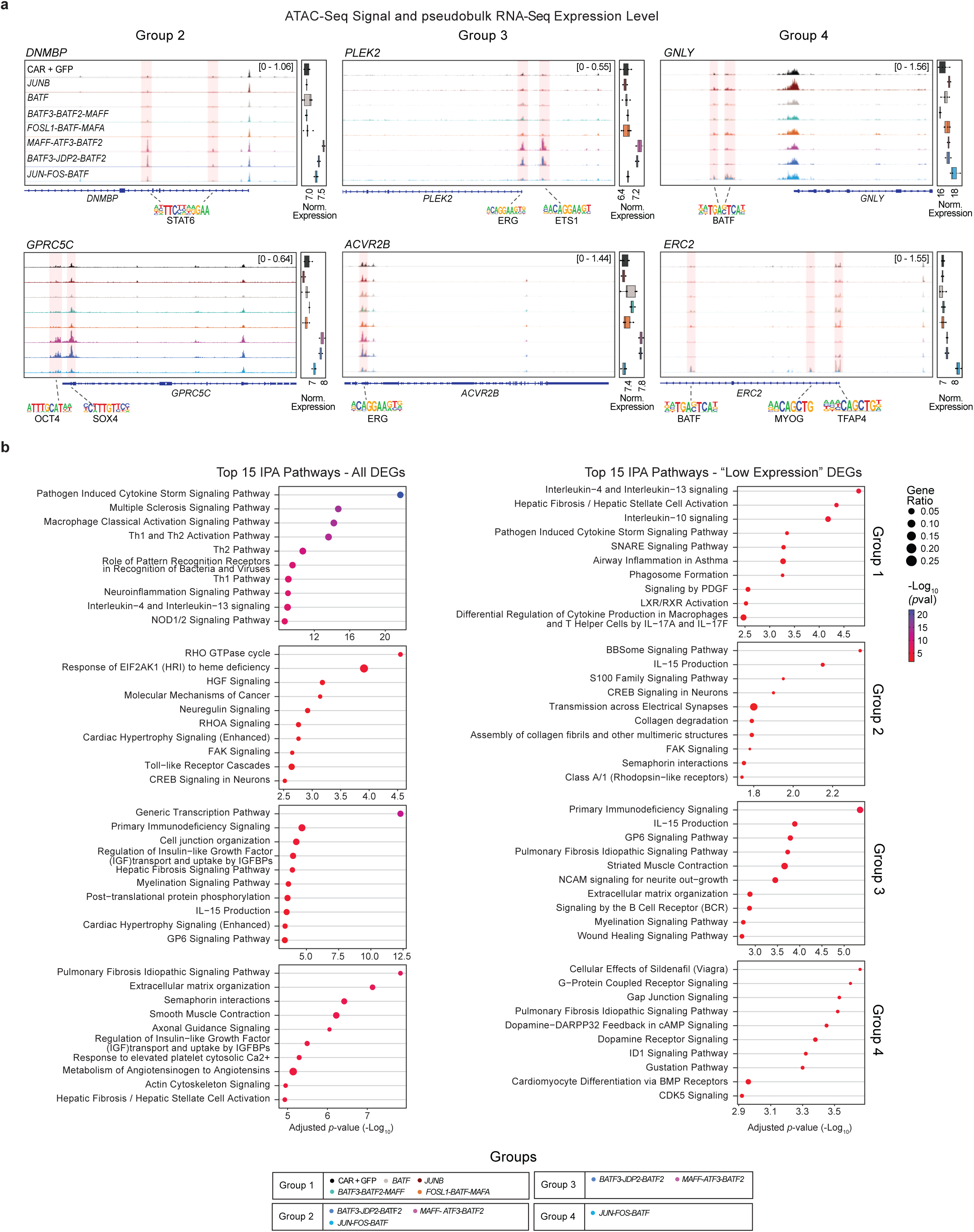
Characterization of DESynR and natural AP-1 TF chromatin accessibility and biological pathway enrichment. **a)** Chromatin accessibility tracks for selected loci along with normalized gene expression (pseudobulk RNA profiles). Average of all human donors (n=3-5) for both data types. **b)** Ingenuity pathway analysis of DEGs (pseudobulk RNA-seq data) from each module. Plots on the left consider all genes and plots on the right consider genes that are in the lowest third of absolute expression in CAR + GFP-control. Top 10 pathways shown. *p*-value from Fisher’s exact test.

